# Pooled sequencing reveals patterns of underlying genomic divergence in *Drosophila* chronotypes

**DOI:** 10.64898/2026.11.14.683300

**Authors:** Arijit Ghosh, Lakshman Abhilash, Vijay Kumar Sharma, KL Nikhil, Vasu Sheeba

## Abstract

Chronotypes, often referred to in the context of sleep timing in humans (early/late sleepers) are heritable variations in timing of circadian behaviours. Chronotype differences among individuals have been associated with a myriad of psychological, metabolic, and physiological dysregulations. Genome-Wide Association Studies (GWAS) in humans have identified several circadian clock gene polymorphisms associated with divergent chronotypes; however, the lists of candidate genes vary across studies based on methodology and sampling strategy thus, necessitating more controlled studies on well-established model systems. Using long-term experimental evolution approach spanning over 20 years, we have derived a unique set of chronotypes in the form of 12 replicate *Drosophila melanogaster* populations that exhibit early/morning and late/evening emergence and have previously reported several circadian clock differences between the populations. We performed pooled DNA sequencing to explore genomic differences underlying early and late chronotypes and identified multiple genomic regions under selection. We observe that the early chronotype populations have primarily accumulated polymorphisms in genes related to photosensitivity, sleep, and the circadian clock, while the late chronotype populations have largely accumulated polymorphisms in genes related to the circadian clock, mRNA splicing mechanisms, and ecdysis. We further predict biological pathways under differential selection in the early and late chronotype populations. We believe these results provide evolutionary mechanisms behind chronotype divergence and will help generate verifiable hypothesis regarding target genes for phase divergence.

## Introduction

Most living organisms ranging from cyanobacteria to modern-day humans exhibit robust daily rhythms in several behaviours and physiological processes. Such 24-h (circadian) rhythms are driven by self-sustaining, endogenous clocks – circadian clocks. Circadian clocks not only drive rhythmic behaviours but also regulate the timing/phase of such behaviours thereby facilitating temporal adaptation of organisms to their environment to maximize fitness (Nikhil and Sharma, 2017). Humans exhibit considerable inter-individual variation in their phase of sleep/wake rhythms. Some are ‘morning people’ with an inherent propensity (not dictated by social norms) to wake up and sleep earlier in the day - ‘early’ chronotypes while ‘late’ chronotypes constitute individuals who inherently wake up and sleep later (Randler et al., 2017). In the modern world, fixed work, school, and social schedules may disproportionately favour certain chronotypes over others leading to a constant misalignment of internal clock with the external time, and this can have obliterating consequences on health. Numerous studies have reported association of chronotype differences with several psychological, metabolic, and physiological dysregulations. For instance, late chronotypes report more frequent psychological and psychosomatic problems (Giannotti et al., 2002; Mecacci and Rocchetti, 1998; Wittmann et al., 2006), daytime tiredness and low night-time sleep quality (Giannotti et al., 2002; Taillard et al., 2003; Volk et al., 1994; Wittmann et al., 2006) compared to early chronotypes. In a recent study the authors found early chronotype students outperformed late chronotypes when school timing was in the morning shift, while late chronotype students performed better in evening classes (Goldin et al., 2020). In another study, where the Seattle school district delayed their secondary school start time by ∼1 hour, there was an increase in median sleep duration by 34 minutes and median grades increased by 4.5% (Dunster et al., 2018). Late chronotype individuals have also more often been associated with being smokers, having lower perceived health, physical inactivity, greater alcohol use, and being more susceptible to sleep interruptions when given early shifts (Juda et al., 2013; Maukonen et al., 2016; Taylor et al., 2020). Thus, there is a need for better understanding of the molecular-genetic underpinnings of chronotype regulation which may potentially enable addressal of several of the associated dysfunctions discussed above.

Studies exploring the genetic basis of chronotypes have reported varying degrees of heritability across human populations (∼50% in the USA, UK, Scandinavia, Brazil, 14% in Hutterites, and 23% in Amazonians, 12-19.4% in UK BioBank data) (Aguiar et al., 1991; Hur et al., 1998; Jones et al., 2016, 2019; Klei et al., 2005; Koskenvuo et al., 2007; Lane et al., 2016; Von Schantz et al., 2015) while others have reported association of chronotypes with various polymorphisms in circadian clock genes such as *CLOCK* (Katzenberg et al., 1998; Mishima et al., 2005), *PER1-3* (Archer et al., 2003; Carpen et al., 2005, 2006; Ebisawa et al., 2001; Pereira et al., 2005), and *ARNTL2* (Parsons et al., 2014). However, some of these reports remain inconclusive as other studies either failed to replicate the findings or observed conflicting or no association of these polymorphisms with chronotypes (An et al., 2014; Barclay et al., 2011; Drake et al., 2015; Iwase et al., 2002; Kunorozva et al., 2012; Osland et al., 2011; Pedrazzoli et al., 2007; Perea et al., 2014; Pereira et al., 2005; Robilliard et al., 2002; Viola et al., 2007). Genome-Wide Association Studies (GWAS) have reported multiple circadian clock genes, genes involved in neuronal signalling, sleep homeostasis and light input pathways to the clock to be associated with sleep chronotypes in humans (Gottlieb et al., 2007; Hu et al., 2016; Jones et al., 2016, 2019; Lane et al., 2016). Interestingly, some of these genes have also been implicated with similar functions in mice (Koike et al., 2012) and *Drosophila* (Zhang and Emery, 2013). While being a powerful strategy, identified candidate genes via GWAS based studies vary across studies for multiple reasons (Kalmbach et al., 2017) thus necessitating further validation of the candidates, which is often not feasible in humans and requires model systems. Moreover, although a vast majority of studies on chronotypes are in the context of timing of human sleep/wake behaviour, studies on other organisms (Aschoff and Wever, 1962; Dominoni et al., 2013; Frías-Lasserre et al., 2019; García-Allegue et al., 1999; Helm and Visser, 2010; Ocampo-Garcés et al., 2006; Refinetti et al., 2016; Schwartz and Smale, 2005; Sheeba et al., 2001; Stuber et al., 2015; Vivanco et al., 2009) highlight that the genetic architecture underlying chronotype/phase regulation is likely to be conserved across species.

To explore the genetic bases of chronotype variation, we turned to a unique resource that was generated using a laboratory-based artificial selection approach (Garland and Rose, 2009). Here 12 populations - 4 sets of genetically independent, large, outbreeding populations of *Drosophila melanogaster* exhibiting early/morning or late/evening emergence chronotypes were derived from their ancestral control populations (henceforth referred to as *early*, *late* and *control* populations Fig. 1A; Kumar et al., 2007, detailed description in methods). The laboratory artificial selection approach has been adopted in a variety of models, from *Escherichia coli* to *Arabidopsis thaliana* and provides numerous advantages over studies on natural populations to explore questions pertaining to adaptation, estimation of population-genetic parameters and genetic architecture of traits as reviewed in Kawecki et al., 2012. Over the course of ∼20 years (∼350 generations) we have reported that the chronotype differences between the *early* and *late* populations may stem from divergent evolution of circadian clock period and amplitude (Kumar et al., 2007, Nikhil, Vaze, et al., 2016) and differential phasing and coupling of the underlying neuronal clocks (Nikhil, Abhilash, et al., 2016), all of which in turn may contribute to differential sensitivity and utilization of light and temperature in the environment (Ghosh et al., 2021; Nikhil et al., 2014; Vaze et al., 2012a, 2012b). These are in accordance with studies on other organisms including humans reporting that chronotypes stem from differences in various clock properties and their interaction with environmental cues (Aschoff and Pohl, 1978; Duffy et al., 2001; Duffy and Wright, 2005; Kerkhof and Van Dongen, 1996; Lehmann et al., 2012; Vivanco et al., 2010). In this report, we leverage these well-characterized *early* and *late* populations that can serve as a suitable system for molecular-genetic exploration of chronotype regulation. To identify putative loci that are associated with chronotype differences we performed pooled-DNA-sequencing of the *early* and *late* populations. Similar approaches using laboratory selection augmented with pooled sequencing strategy has successfully aided exploration of genetic bases of longevity and aging (Burke et al., 2010; Remolina et al., 2012), body size (Turner et al., 2011), hypoxia tolerance (Zhou et al., 2011), courtship song (Turner and Miller, 2012), bristle development (Cassidy et al., 2013), adaptation to novel environments (Orozco-Terwengel et al., 2012), temperature (Tobler et al., 2014), and diet (Reed et al., 2014) in *D. melanogaster*.

**Figure 1.**
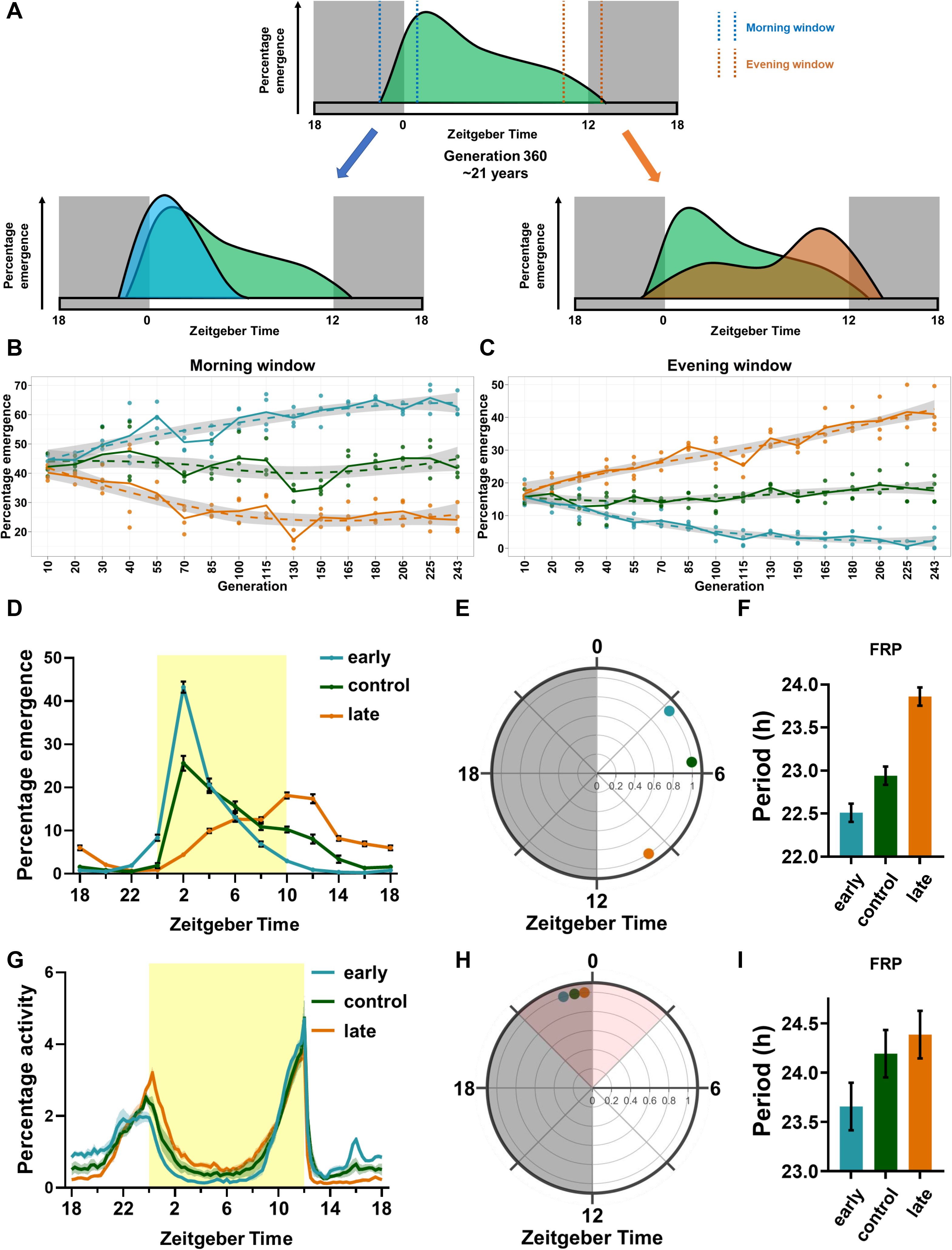
Long-term laboratory selection produces divergent emergence and activity chronotypes. (A) Schematic of the selection regime, showing morning and evening collection windows and the resulting early and late emergence profiles. (B, C) Percentage emergence within the morning (B) and evening (C) windows across generations of selection. Each small point represents one replicate population. Solid lines connect means across the four replicate populations at each generation; dashed lines show LOESS-smoothed trends, with shading indicating 95% confidence intervals for the LOESS fits. (D) Emergence profiles under a 12 h light:12 h dark cycle. (E) Polar representation of emergence timing. (F) Free-running period (FRP) of the emergence rhythm. (G) Locomotor activity profiles of males under a 12 h light:12 h dark cycle. (H) Polar representation of activity timing. (I) FRP of the locomotor activity rhythm. In E and H, angle indicates mean phase and radial distance indicates vector strength. Measurements in B–I are based on four independent replicate populations per selection regime (n = 4). Emergence measurements in B–F pooled males and females; locomotor activity measurements in G–I used males only. Error bars and shaded bands in D and G indicate the standard error of the mean (SEM). Bars in F and I show mean free-running periods with 95% confidence intervals. Statistical comparisons of free-running periods used randomized-block ANOVA and Tukey’s multiple-comparisons test. Blue indicates early populations, green control populations, and orange late populations. ZT, Zeitgeber time; ZT0 denotes lights on.

Here we present evidence that in the *early* and *late* populations, in addition to genomic regions undergoing positive selection, the two populations also accumulated differential enrichment of single Nucleotide Polymorphisms (SNPs). Specifically, we found that genes involved in photosensitivity, sleep, and circadian clocks in *early* populations, and mRNA splicing, ecdysis and circadian clocks in *late* populations to have accumulated more SNPs compared to the *control* flies. We further predict different pathways plausibly under differential selection in *early* (circadian rhythm, notch signaling, spliceosome, metamorphosis and nervous system processes) and *late* (mRNA splice site selection, spliceosome, response to stimulus, phototransduction, circadian rhythm and metabolic pathways) populations. Remarkably our study identifies such differences consistently across four genetically independent replicate populations further emphasizing the robustness of the identified putative loci and thus pave the way for future studies aimed at exploring the genomic bases of chronotype regulation.

## Materials and methods

### Laboratory selection protocol to derive *early* and *late* populations

To initiate this project, four sets of genetically independent *Drosophila melanogaster* populations were used to artificially select for morning eclosion (*early* populations) and evening eclosion (*late* populations) timing (Kumar et al 2007). At each generation, ∼300 eggs are collected and placed in glass vials which are maintained in a light-proof, temperature-controlled cubicle under a light-dark cycle of 12:12 h-(LD12:12), 25°C ± 0.5°C, and 65% ± 5% RH. Flies emerging from ZT21-ZT01 (ZT0 is Zeitgeber Time 0, when light comes ON) are collected to form the breeding pool for the next generation of *early* flies, while flies emerging from ZT9-ZT13 form the breeding pool for the next generation of *late* flies (henceforth, denoted as *early_1-4_*and *late_1-4_*). In contrast, for the four *control* populations, flies emerging throughout the day are collected to make up the breeding pool for the next generation (henceforth*, control_1-4_*). This collection is carried out for 3-4 days and a total ∼1200 adult flies with ∼1:1 sex ratio of each of the 12 populations are maintained in Plexiglas™ cages (25 cm×20 cm×15 cm) with petri plates with banana-jaggery culture media. The flies are maintained on a 21 day discrete generation cycle, and all experiments were done with progeny of flies that experienced one generation of common rearing (standardized) to avoid confounding factors due to maternal effects (Bonduriansky and Day, 2009). For more details of the selection regime, see Kumar et al. (2007), and Abhilash et al. (2019). All experiments described in this report were done between generations 260 to 310.

### DNA sample preparation, sequencing, and variant identification

Approximately 500 individuals (250 males and females each; 4-5 days old) were randomly sampled from each population (total 12 samples with *early_1-4_*, *control_1-4_*, and *late_1-4_*) at generation 260. Total genomic DNA was extracted with the DNeasy Blood and Tissue Kit (#69504, QIAGEN, MD, USA), subjected to quality check (Nanodrop QC and Qubit QC) and paired-end libraries of each sample were prepared for sequencing with tags (Illumina HiSeq manufacturer’s protocol). The Illumina HiSeq generated 150 paired-end reads (∼80-100 million per sample) which were quality checked using FastQC (Andrews, 2010). These raw reads were processed by Cutadapt (Martin, 2011) to remove adapters and trim low-quality bases. A ∼100X depth of coverage was aimed for as this was a pooled-sequencing experiment for reliable variant identification (Futschik and Schlötterer, 2010). The reads from all populations were then aligned to the D. melanogaster reference genome (BDGP6) using default parameters for Bowtie 2 (Langmead and Salzberg, 2012).

SNPs and Insertion or Deletions (INDELs) were identified with SAMtools1.2 and BCFtools1.2 (Li et al., 2009) using a read depth threshold ≥ 20 and mapping quality threshold ≥ 30. mpileup files were generated using SAMtools1.2 and further processed using VarScan2 (Koboldt et al., 2009) for identification of population specific markers between the samples in a group. VCF files were created for each of the 12 samples with BCFtools1.2 and both VCF and mpileup files were used for downstream analyses. We only analyzed SNPs, because they were more numerous and partly because INDEL analyses are not standardized for pooled sequencing studies.

### Calculation of Tajima’s Pi (π), Tajima’s D, and Watterson’s Theta (θ)

Population genomic parameters (Tajima’s Pi (π), Tajima’s D, and Watterson’s Theta (θ_w_)) were calculated using PoPoolation_1.2.2 (Kofler, Orozco-terWengel, et al., 2011) and the input mpileup files for each sample. π is a measure of DNA sequence variation based on the average pairwise distance between all samples and θ is a measure of DNA sequence variation based on observed number of segregating sites (polymorphic sites), and numbers of chromosomes per sample. *D* is a statistic that measures the difference between estimators, θ*_w_* (expected heterozygosity) and π (observed heterozygosity) (Biswas and Akey, 2006; Tajima, 1989). Positive values of *D* arise from an excess of intermediate frequency alleles (higher heterozygosity than expected) and can result from population bottlenecks, structure and/or balancing selection. Negative values of *D* indicate an excess of low frequency alleles (lower heterozygosity than expected) and can result from population expansions or positive selection (Biswas and Akey, 2006). Discovery of regions with significantly low *D* was determined using the Two-stage linear step-up procedure of Benjamini, Krieger and Yekutieli, with Q = 5% using GraphPad Prism 8 (13213 independent t tests for each comparison). Since π, *D*, and θ are sensitive to variation in coverage, reads were subsampled to a uniform coverage of 20 for removing any bias in calculations before we calculated π, *D*, and θ*_w_* for 50Kb windows with a 10Kb step size using the *Variance-sliding.pl* script that is corrected for pooled-sequencing (Kofler, Orozco-terWengel, et al., 2011).

### Allele frequency-based differentiation tests

We used the sorted bam files to create pairwise mpileup files – one mpileup file for *control_1-4_* and *early_1-4_*, and another mpileup file for *control_1-4_* and *late_1-4_*. These two mpileup files were further converted into PoPoolation2 compatible SYNC files using the Java program mpileup2sync.jar setting a minimum quality of 30. These two SYNC files were then used to calculate pairwise F_ST_ values between each SNP of matched populations (*control_1_* vs *early_1_*, *control_1_* vs *late_1_*, *control_2_* vs *early_2_*, *control*_2_ vs *late_2_* and so on). F_ST_ is a statistic that quantifies levels of differentiation between subpopulations and can be used to identify SNPs whose allele frequency varies significantly between two subpopulations (Hartl and Clark, 1989). We applied a Z-transformation on the F_ST_ values from four different replicates of *control-early* and *control-late* comparisons and used a 5% threshold using the *quantile* function in R (version 4.0.2) to identify significant SNPs (R Core Team, 2020). All significantly differentiated SNPs common among all four replicates for each pairwise comparison were considered for further analyses. Only SNPs from all replicates were considered for further analyses because SNPs changing in allele frequency in all four replicates are much more likely to be due to the similar selection pressure they experience, than just by genetic drift, which might have been acting differentially in different replicate populations. Next, we performed a Cochran-Mantel-Haenszel (CMH) test for repeated tests of independence for every SNP. The CMH test is used for detecting significant and consistent changes in allele frequencies, when independent measurements of allele frequencies are available, as in our case where we can treat the replicate populations as genetically independent samples. CMH test was implemented by the *cmh-test.pl* code with *min-count* 2 per population, *min-coverage* of 30, and *max-coverage* of 400 between *control-early* and *control-late* (Kofler, Pandey, et al., 2011). The traditional CMH tests are known to be less conservative and may give rise to numerous false positives (Wiberg et al., 2017). We took a permutation based approach to determine significance thresholds for respective CMH tests (Graves et al., 2017). We used permutation to generate null distributions, and then set a genome wide significance threshold based on the smallest *p* values in that distribution (1000 permutations, 5% cut-off). Using this strict cut-off, we identified 333 SNPs in *early* populations (CMH*p_corrected_*= 1.396817e-58) and only 3 SNPs in *late* populations (CMH*p_corrected_*= 4.556171e-55) where allele frequencies changed significantly from *control* populations. Therefore, we lowered the cut-off for *late* populations (CMH*p_corrected_* = 3.046445e-27) to include SNP numbers similar to that of the *early* populations by choosing top 333 SNPs with lowest CMH*p* value. Manhattan plots of results from pairwise F_ST_ comparisons and CMH tests were plotted using the CMplot (LiLin-Yin, 2020) package in R (version 4.0.2) (R Core Team, 2020).

### Identification of genome-wide selective sweeps

We took a more direct approach to identify genomic regions that have likely undergone selective sweeps in the *early*, *control*, and *late* populations using *Pool-hmm* (Boitard et al., 2013). We ran *Pool-hmm* for each chromosomal arm separately with respective populations size for pooling individuals with *k* value of 1^-10^, and θ value as calculated before for each sample (θ*_w_*). Further data processing was done using the data.table package (Dowle and Srinivasan, 2020) and plotted using the karyoploteR package (Gel, 2020; Gel and Serra, 2017) in R (version 4.0.2) (R Core Team, 2020).

### Analysis of identified variants

After identifying the significantly differentiated variants in the *control-early* and *control-late* comparisons, we segregated locations common to both *early* and *late* populations which have diverged from the *controls*. Thus, we ended up with three exclusive groups of SNPs, *early* exclusive, *late* exclusive, and *early-late* overlap. We were interested in the *early* and *late* exclusive group of variants and proceeded to annotate them using SnpEff (version 4.3t) variant effect predictor (Cingolani et al., 2012). SNPs were categorized based on estimation of their putative impact/deleteriousness (low, moderate, high, and modifier) and their effect type-description of consequence (see results section). Gene ontology (GO) analysis and pathway enrichment analysis was carried out with the identified genes from SnpEff using g:Profiler (Reimand et al., 2016). Further manual categorization of different GO terms was done. To identify statistical enrichment of overlap between significantly differentiated SNPs harboring genes among different sets (Tajima’s D comparison, pairwise FST comparison, and CMH tests), we used the SuperExactTest package (Wang et al., 2022) in R (version 4.0.2) (R Core Team, 2020). Protein-protein interaction networks were constructed using STRING (https://string-db.org/) (Szklarczyk et al., 2021), and later analyzed and visualized using StringApp (Doncheva et al., 2019) in Cytoscape 3.7.2 (Shannon et al., 2003). Tajima’s *D*, π, θ_w_, KEGG enrichment analysis, and GO enrichment analysis results were plotted with ggplot2 package (Wickham, 2016; Wickham et al., 2020) in R (version 4.0.2) (R Core Team, 2020).

## Results

### Long term laboratory selection for morning and evening emergence has resulted in divergent circadian phenotypes in *early and late* populations

Eclosion/emergence - the act of a developmentally complete fly emerging from its puparium is a phenomenon exhibited by many insects. Emergence in *D. melanogaster* is regulated by circadian clocks and consequently we observe circadian rhythms in emergence at a population level where majority of flies emerge in the early morning while very few to no flies emerge in the later hours of the day through the night (*Fig. 1A*). While mutations in circadian clock genes can advance or delay the phase/timing of emergence thus rendering them early-like and late-like chronotypes, a long-term (> 20 years) laboratory selection protocol has produced four sets of *D. melanogaster* populations exhibiting morning/early and evening/late emergence (see Materials and Methods; *Fig. 1A*). As a result of several generations of selection, the phase of emergence (emergence chronotypes) has diverged between the populations with the proportion of flies emerging in the morning window increasing significantly in the *early* populations while that of evening window increased for the *late* populations (*Fig. 1B–F;* Kumar et al., 2007). Over the years, we have shown that the *early* and *late* populations also exhibit similar chronotype differences in activity-rest rhythms (Fig. 1G,H; Abhilash et al., 2020), different circadian clock periods (Fig. 1F,I; Nikhil, Abhilash, et al., 2016) and sensitivity to environmental stimulus (Ghosh et al 2021).

### Different genomic regions in the *early* and *late* populations show signatures of selection

To identify *early* and *late* populations-specific genomic regions that had diverged from the *control* populations, we first identified SNPs passing all quality controls, calculated different nucleotide diversity and heterozygosity matrices, and then employed a broad range of allele frequency-based methods to identify genomic regions under differential selection in the *early* and *late* populations compared to *control*. We identified a total of 1.37 million SNPs spanning 17477 genes in the *early* populations and 1.43 million SNPs spanning 17515 genes in *late* populations when compared to the *control* populations (*Fig. 2A,B*). To ensure that we increase the chances of considering only allele frequency changes due to selection and not due to other forces such as genetic drift, we selected only the SNPs that passed the strict quality control of presence in all four replicate populations. The SNPs did not show any discernible spatial pattern of distribution or bias in SNP calling and were distributed all over the five major chromosomal arms (*Fig. 2A,B*). The SNP density pattern across chromosomal arms were similar for both *early* and *late* populations, and in both sets of populations, the X chromosome had lower number of SNPs compared to the other arms (*Fig. 2A,B*). There were many more variants identified in the mpileup files compared to the 1.37 and 1.43 million SNPs used for *early* and *late* populations respectively, but while using the mpileup files for different population genomic parameter estimations (π, *D*, and θ*_w_*), we kept similar quality control of SNPs, so that similar number of SNPs are used for these estimations (Fig. S1 and S2).

**Figure 2.**
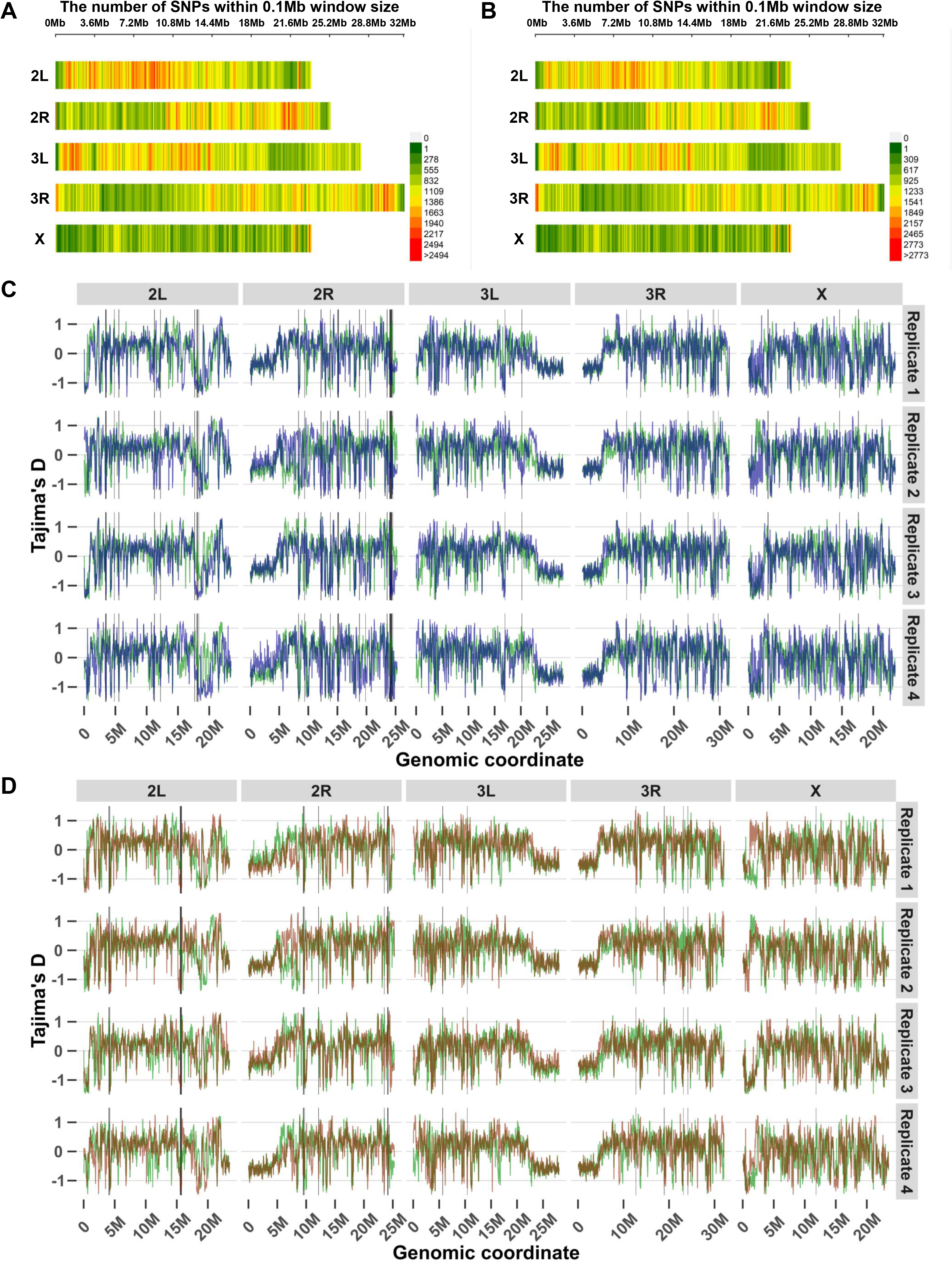
Genome-wide SNP density and Tajima’s D. (A, B) SNP counts in non-overlapping 0.1 Mb windows across five major chromosome arms in the control-early (A) and control-late (B) comparison datasets. Approximately 1.37 million and 1.43 million SNPs, respectively, passed quality control across the four replicate populations. Color scales indicate the number of SNPs per window. (C, D) Tajima’s D in 50 kb windows advanced in 10 kb steps for control and early (C), and control and late (D), populations. Columns indicate chromosome arms and rows indicate the four replicate populations. Blue indicates early, green control, and red late populations. Genome-wide median D values were 0.121 for control, 0.114 for early, and 0.138 for late populations.

We calculated Tajima’s D (*D*) for 50Kb windows with a step size of 10Kb to identify genomic regions significantly under selection (regions with significantly lower negative *D* values compared to *control* populations) in the *early* and *late* populations. All 12 populations showed overall high levels of heterozygosity and standing genetic variations (measured by *D* and π) as expected from large, outbreeding, random mating populations such as these, but overall *D* values were higher than were previously reported for different natural populations (Croze et al., 2017; Fabian et al., 2012). Median genome-wide *D* value of the respective populations were – *control*: 0.121 (range: −1.333 to 1.233), *early*: 0.114 (range: −1.393 to 1.192), and *late*: 0.138 (range: −1.39 to 1.17). We identified 159 genomic regions of 10Kb intervals (total 1.59 Mb region) in *early* populations and 87 regions in *late* populations with significantly lower *D* values than the *control* populations indicating that these regions are under stronger positive selection in these regions (Fig. 2C,D). In the *early* populations, these regions identified to be under positive selection harbor 296 genes (including pseudogenes, protein coding genes, miRNA, snoRNA, lncRNA, tRNA, and antisense lncRNA genes) while those in the *late* populations harbor 112 genes. Some noteworthy genes within these regions are listed in *Table 1*.

**Table 1.** Selected genes in regions identified by Tajima’s D comparisons. Selected genes within genomic regions with lower Tajima’s D in early or late populations than in control populations. Functions and references describe reported biological roles and do not establish causal effects of the variants on chronotype.

| Population | Gene name | Function | Reference |
|---|---|---|---|
| early | nord | blue light induced gene | Hall et al., 2018 |
| early | Arrestin 1 | phototransduction | Damulewicz et al., 2019 |
| early | nervy | regulates PDF expression | Duvall and Taghert, 2012 |
| early | Thor | Contributes to circadian functions | Nagoshi et al., 2010 |
| early | reduce ocelli | ocelli and photoreceptor formation | Caldwell et al., 2007; Vosshall and Young, 1995 |
| early | pickpocket 29 | neurotransmission | Hill et al., 2017 |
| early | mAChR-A | regulates light regulate calcium response in LNVs | Qin et al., 2019 |
| early | spaghetti | regulator of DBT | Means et al., 2015 |
| early | Leucyl-tRNA synthetase | blue light induced gene | Hall et al., 2018 |
| early | dunce | mutants show augmented light-induced phase delay and shortened period | Levine et al., 1994 |
| early | ninaD | rhodopsin biosynthesis | Gu et al., 2004 |
| early | timeless | core circadian clock gene | Sehgal et al., 1994 |
| late | Blimp-1 | pupation timing, fat body timer | Akagi et al., 2016 |
| late | Rootletin | sensory perception of sound and touch | Chen et al., 2015 |
| late | Tsc1 | keeps circadian period close to 24 hours and maintains robust circadian rhythms | Zheng and Sehgal, 2010 |
| late | spaghetti | regulator of DBT | Means et al., 2015 |

We also calculated π (average pairwise observed nucleotide diversity) in 50Kb windows with a step size of 10Kb along all the five major chromosomal arms of all populations (Fig. S1). Median π values were 0.00262 (range: 0-0.01; SD: 0.0018) for *control*, 0.00242 (range: 0-0.12; SD: 0.001772) for *early*, and 0.002641 (range: 0-0.011; SD: 0.001798) for *late* populations, reaffirming the high amount of heterozygosity and standing genetic variation within each population.

These analyses suggest that our populations had high standing genetic variation within them, and despite undergoing selection for ∼260 generations, the *early* and *late* populations have maintained genome-wide heterozygosity and have undergone significantly more positive selection compared to the *control* population at multiple genomic regions (Fig. 2C,D; Figs. S1 and S2).

### Identification and characterization of significant SNPs by quantifying differential allele frequency changes between populations

While Tajima’s D (*D*) values are useful for investigating effect of selection of each population separately, comparing allele frequencies of different populations with each other is necessary for identifying loci undergoing significant allele frequency changes in the *early* and *late* populations as compared to the *control* population. We next investigated the patterns of genetic differentiation in *early* and *late* populations in response to the imposed selection by calculating pairwise F_ST_ between all SNPs in *control-early* and *control-late* population pairs. In the four replicates analyzed, median autosomal F_ST_ values in *early* population varied from 0.015 to 0.024, and in X chromosome, F_ST_ varied from 0.017 to 0.023 (Fig. 3A). In the case of *late* populations, median autosomal F_ST_ values among four replicates varied from 0.013 to 0.016, and in X chromosome it varied from 0.013 to 0.017 (Fig. 3B). Owing to its smaller effective population size, the X chromosome has higher average F_ST_ values than the autosomes as shown previously (Hutter et al., 2007; Kapun et al., 2020) as also reflected in our populations. Overall, the four replicates showed different degrees of differentiation reaching a range of 0 to 1 in replicate 2 of *early* populations, while in other three replicates, it varied from 0 to 0.95, 0.88, and 0.93. However, in *late* populations, F_ST_ values varied from 0 to 0.67, 0.78, 0.64, and 0.83 (Fig. 3B). Next, we focused on loci where F_ST_ values in all four replicates show high values. We achieved this by calculating zF_ST_ values (a Z-transformation on the pairwise F_ST_ values of *control-early* and *control-late* comparisons) and using a 95-percentile cut-off for each replicate and choosing only those loci where zF_ST_ values pass the cut-off in all four replicates (Fig. 3A,B). We consequently identified 686 SNPs in the *early* populations (maxF_ST_ =1, minF_ST_ = 0.15, maxzF_ST_ = 14.45, minzF_ST_ = 1.98) and 79 SNPs in *late* populations (maxF_ST_ = 0.65, minF_ST_ = 0.12, maxzF_ST_ = 10.12, minzF_ST_ = 2.06) with allele frequencies significantly diverged from *control* populations in all four replicates. Although there is no consensus about cutoffs of F_ST_ values for significant differentiation, Hartl and Clark (1989) proposed a simple classification system, where F_ST_ <0.05 indicates “little genetic differentiation”, 0.05-0.15 = “moderate genetic differentiation”, 0.15-0.25 = “great genetic differentiation”, and >0.25 = “very great genetic differentiation”. While we adhere to our zF_ST_ based 95-percentile cut-offs, we note most of our identified significant SNPs have F_ST_ values greater than 0.15 (*early* average 0.361; range: 0.15 to 1, *late* average 0.229; range: 0.12 to 0.65), which can be categorized as “great to very great” genetic differentiation. Among the 686 SNPs identified in the *early* populations spanning over 127 genes, SnpEff predicted 112 to have an estimated “low” effect of putative impact/deleteriousness, 15 “moderate” effect SNPs, and 559 “modifier” effect SNPs (Supplementary Table S1; Fig. 3C,D). Among the 79 SNPs identified in the *late* populations spanning over 47 genes, SnpEff predicted 9 to have “low” effect of putative impact/deleteriousness, 2 “moderate” effect SNPs, and 68 “modifier” effect SNPs (Supplementary Table S2; Fig. 3C,D). From the F_ST_ analysis, the genes directly or indirectly related to circadian or other functions identified in *early* and *late* flies are listed in Table 2. Interestingly, there was no overlap between the loci identified in *early* and *late* populations. In *early* populations, gene ontology (GO) terms represented in the summaries included “biological regulation”, “regulation of signaling”, “regulation of signal transduction”, “nervous system processes’, “instar larval or pupal development”, “metamorphosis” (Fig. S3A) while those for the *late* populations included “localization”, “primary metabolic process”, “response to stimulus” and “transport” (Fig. S3B). Many identified genes harbour multiple differentiated SNPs; the 40 genes or annotated loci with the largest counts in each comparison are shown in Fig. 5A,B. The predicted consequence distributions are also summarized in Fig. S5A,B.

**Figure 3.**
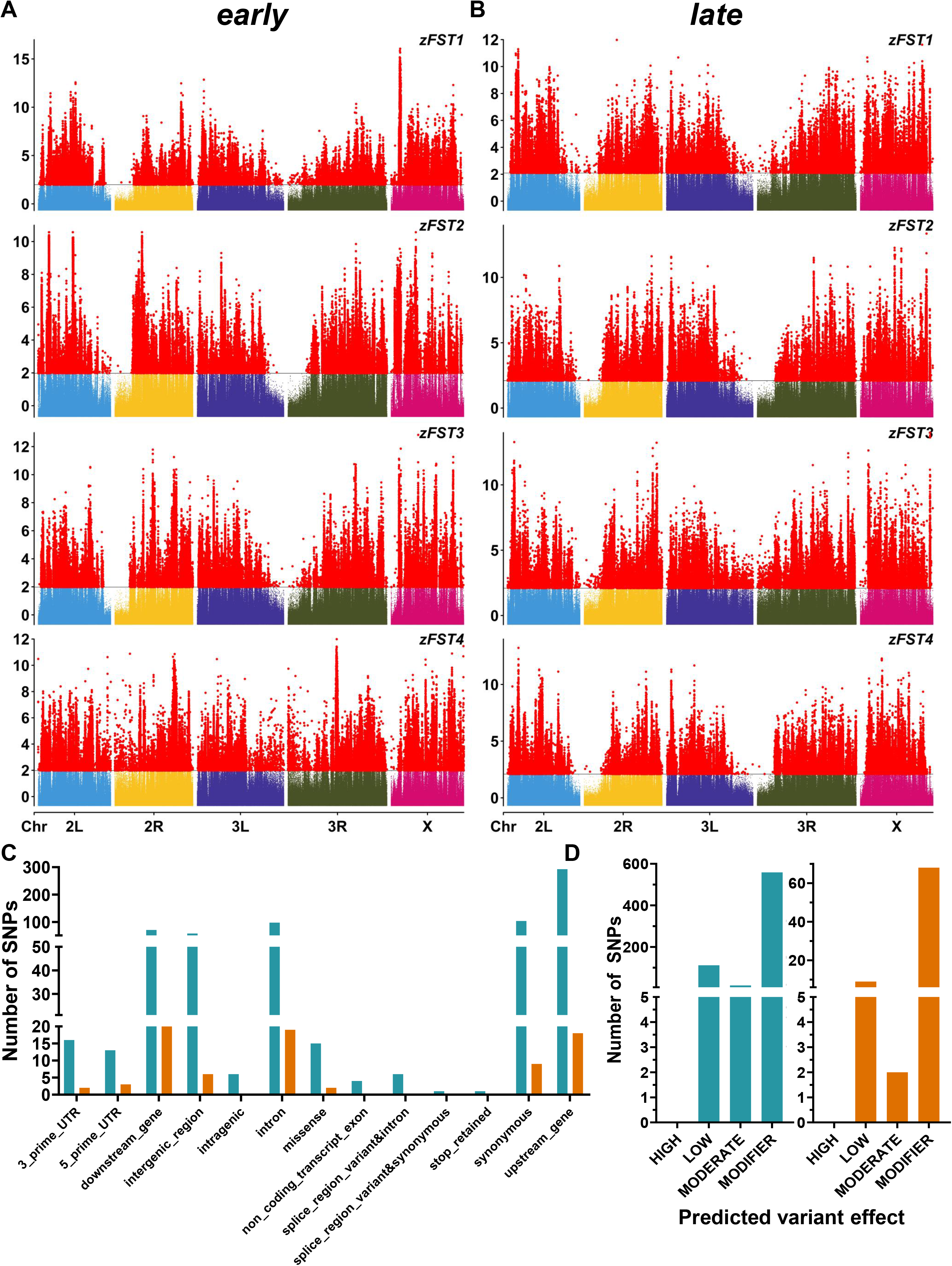
Allele-frequency differentiation and predicted variant consequences from FST comparisons. (A, B) Manhattan plots of z-transformed FST (zFST) across five major chromosome arms for the four matched control-early (A) and control-late (B) replicate comparisons. Red points mark SNPs exceeding the replicate-specific 95th-percentile cutoff; chromosome colors identify the remaining points. SNPs exceeding the cutoff in all four replicates were retained, and positions shared between the early and late comparisons were removed to obtain early-exclusive and late-exclusive sets. (C) Predicted sequence-consequence categories for the 686 early-exclusive and 79 late-exclusive SNPs. (D) Corresponding SnpEff predicted-impact categories. Teal indicates early-exclusive SNPs and orange late-exclusive SNPs. Broken vertical axes permit categories with different counts to be displayed together. Predicted impact is an annotation category, rather than an experimentally measured effect.

**Table 2.** Selected genes identified by pairwise FST comparisons. Selected genes associated with SNPs passing the 95th-percentile zFST threshold in all four matched early–control or late–control replicate comparisons. Functions and references describe reported biological roles.

| Population | Gene name | Function | Reference |
|---|---|---|---|
| early | kirre | sleep traits | Harbison et al., 2017 |
| early | vrille | core clock component | Panda et al., 2002 |
| early | notch | compound eye development | Kahali et al., 2009 |
| early | timeless | core clock component | Sehgal et al., 1994 |
| early | Egfr | consolidation and maintenance of sleep | Foltenyi et al., 2007 |
| early | Ir64a | odorant receptor | Ai et al., 2013 |
| early | nompC | response to auditory stimulus and startle response | Boyd-Gibbins et al., 2021; Zhang et al., 2013 |
| early | dpr8 | synapse organization | Carrillo et al., 2015 |
| early | Nlg4 | sleep, expressed in clock neurons | Li et al., 2013 |
| early | Fbxl4 | negative regulation of circadian sleep/wake cycle | Li et al., 2017 |
| early | bruchpilot | adult locomotory behavior | Wagh et al., 2006 |
| early | Basigin | photoreceptor cell morphogenesis | Munro et al., 2010 |
| early | CCAP-R | neuropeptide signaling pathway, ecdysis | Cazzamali et al., 2003 |
| early | Eip93F | adult determinant during fly morphogenesis | FlyBase |
| early | Tyramine β hydroxylase | sleep | Crocker and Sehgal, 2008 |
| early | Thor | contributes to circadian functions | Nagoshi et al., 2010 |
| early | sgg | modulates periodicity of the circadian clock | Martinek et al., 2001 |
| early | 5-HT1B | circadian photosensitivity | Yuan et al., 2005 |
| late | Heterochromatin protein 5 | gene silencing by heterochromatin | FlyBase |
| late | unplugged | brain development | Hirth et al., 2003 |
| late | Cyp4p1, Cyp4p2, Cyp4p3 | metabolism of insect hormones | FlyBase |
| late | Netrin-A | compound eye development | Akin and Zipursky, 2016 |
| late | Rootletin | sensory perception of sound and touch | Chen et al., 2015 |
| late | Sirt4 | determination of adult lifespan | Wood et al., 2018 |

Furthermore, to understand if specific biological pathways have been under differential selection in *early* and *late* populations, we performed a pathway enrichment analysis using g:Profiler (Reimand et al., 2016). The KEGG pathway summaries for *early* populations included “metabolic pathways”, “circadian rhythm - fly”, “apoptosis - fly”, “notch signaling pathway”, “longevity regulating pathway – multiple species”, “spliceosome”, “drug metabolism – other enzymes”, whereas in *late* populations, the pathway summaries included “metabolic pathways”, “nicotinate and nicotinamide metabolism”, “longevity regulating pathway – multiple species”, “FoxO signaling pathway”, “phagosome”, “lysosome” (Fig. 6A,B; gene and locus counts are shown in Fig. 5A,B).

We used the CMH test to identify loci where allele frequency has concurrently changed among all four replicate populations of *early* and *late* populations compared to that of *control* populations. In *early* and *late* populations, we find 287 exclusive SNPs significantly different from *control* populations using the cut-offs described in materials and methods (Fig. 4A,B). We also identify 46 mutual SNPs in *early* and *late* populations which have significantly different allele frequency compared to control populations. In the *early* populations, these 287 SNPs span over 65 genes. Among these SNPs, SnpEff predicted 115 to have an estimated “low” effect of putative impact/deleteriousness, 24 “moderate” effect SNPs, 147 “modifier” effect, and 1 “high” effect SNPs (Supplementary Table S3; Fig. 4C,D). In the *late* populations, these 287 SNPs span over 107 genes. Genes directly or indirectly related to circadian or other functions identified in *early* and *late* flies identified by the above method are listed in *Table 3*. Among these SNPs, SnpEff predicted 110 to have an estimated “low” effect of putative impact/deleteriousness, 27 “moderate” effect SNPs, and 150 “modifier” effect SNPs (Supplementary Table S4; Fig. 4C,D). Among the 46 SNPs spanning over 10 genes, which were common between *early* and *late* populations, SnpEff predicted 31 to have an estimated “low” effect of putative impact/deleteriousness, 4 “moderate” effect SNPs, and 11 “modifier” effect SNPs. The 46 shared SNPs comprised 4 upstream-gene variants, 31 synonymous variants, 4 intron variants, 1 downstream-gene variant, 2 3′-UTR variants, and 4 missense variants of moderate predicted impact (Supplementary Table S9). *timeless* (core clock component) is one of the major genes which accumulated 14 SNPs common to both *early* and *late* populations. In *early* populations, GO terms represented in the summaries included “transcytosis”, “RNA polymerase III type 2 promoter sequence-specific DNA binding”, “Mre11 complex”, “transcription factor binding”, and the KEGG pathway summaries included “metabolic pathways”, “circadian rhythm - fly”, “notch signaling pathway”, “dorso-ventral axis formation”, “longevity regulating pathway – multiple species”, “lysosome” (Fig. 6C; Fig. S3C). In *late* populations, GO terms represented in the summaries included “miRNA loading onto RISC involved in gene silencing by miRNA”, “chemoattractant activity”, “mRNA splice site selection”, “sensory perception of sound”, “sensory perception of mechanical stimulus”, and the KEGG pathway summaries included “metabolic pathways”, “longevity regulating pathway – multiple species”, “circadian rhythm - fly”, “retinol metabolism”, “phototransduction - fly”, “FoxO signaling pathway”, “citrate cycle (TCA cycle)”, “ubiquitin mediated proteolysis”, “spliceosome”, “drug metabolism – other enzymes” (Fig. 6D; Fig. S3D). The top 40 genes or annotated loci by differentiated-SNP count in the CMH comparisons are shown in Fig. 5C,D, and the predicted consequence distributions in Fig. S5C,D. At the strict permutation-derived threshold, only three *late*-comparison SNPs were retained (Fig. S3E); the matched top-333 *late* set uses the explicitly relaxed threshold described in Methods.

**Figure 4.**
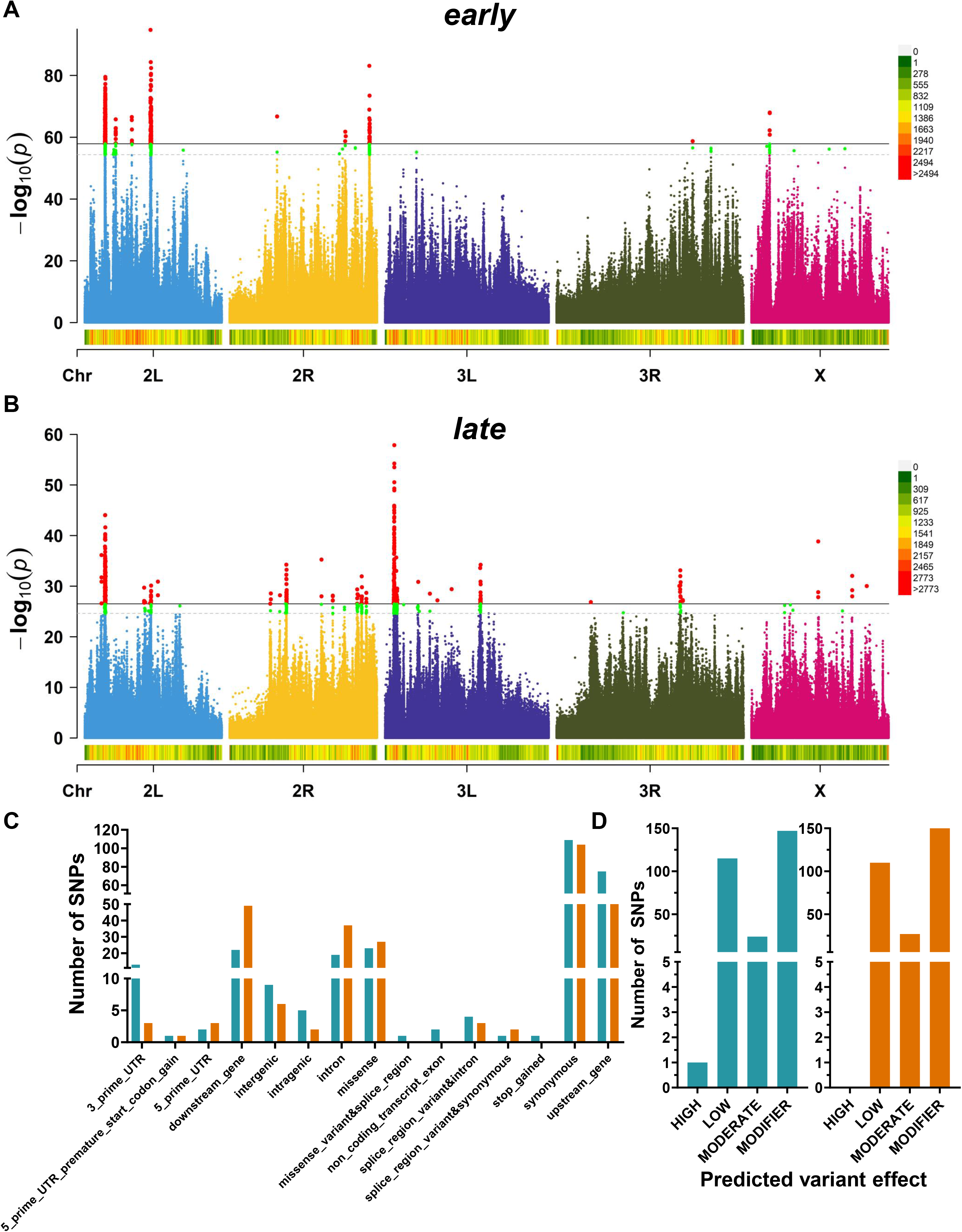
Allele-frequency differentiation and predicted variant consequences from Cochran-Mantel-Haenszel (CMH) tests. (A, B) Manhattan plots of -log10(p) across five major chromosome arms for control-early (A) and control-late (B) comparisons. The upper solid lines delimit the top 333 SNPs (red points). For the early comparison, the permutation-derived 5% cutoff was p = 1.396817 × 10; the late cutoff was relaxed to p = 3.046445 × 10 ² to retain the same number of SNPs. The lower dashed lines delimit the top 551 SNPs, with the additional SNPs shown in green; the early cutoff was obtained from the 10% permutation screen and the late set was matched in size. The top-551 sets are shown for comparison and were not used for the annotations in C and D. SNP counts in non-overlapping 0.1 Mb windows are shown beneath each plot, with color scales at right. (C) Predicted sequence-consequence categories for early-exclusive and late-exclusive SNPs from the top-333 sets. (D) Corresponding SnpEff predicted-impact categories. After removing 46 shared positions, each exclusive set contains 287 SNPs. Teal indicates early-exclusive SNPs and orange late-exclusive SNPs. Broken vertical axes permit categories with different counts to be displayed together. The relaxed late set is a rank-matched candidate set, not a set established by the original permutation significance threshold.

**Figure 5.**
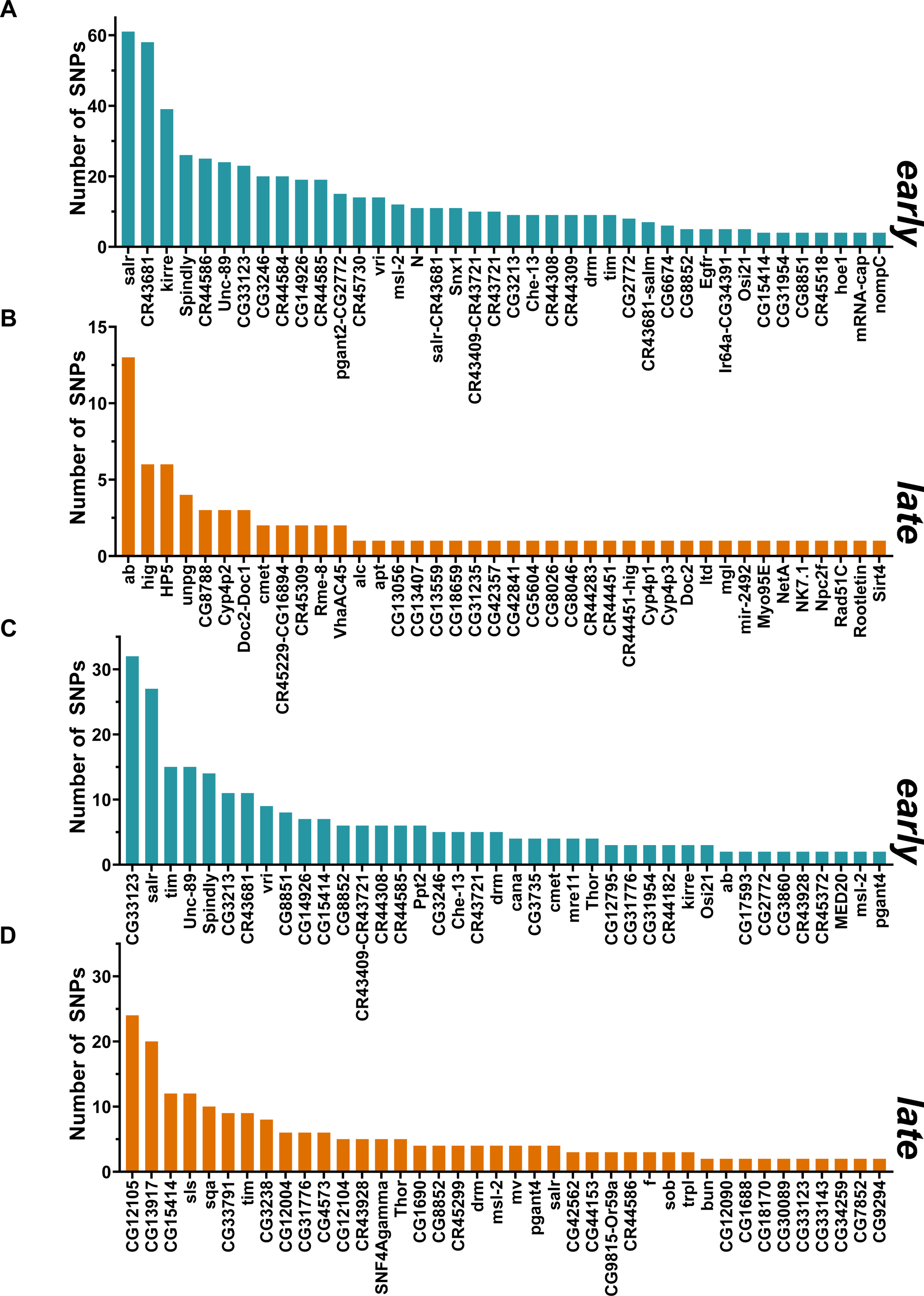
Annotated genes and loci with the largest numbers of differentiated SNPs. (A, B) SNP counts for the top 40 annotated gene or locus labels in the early-exclusive (A) and late-exclusive (B) FST sets. (C, D) Corresponding counts in the early-exclusive (C) and late-exclusive (D) CMH sets. Teal indicates early and orange late populations. Compound labels denote the supplied annotations for intergenic loci.

**Figure 6.**
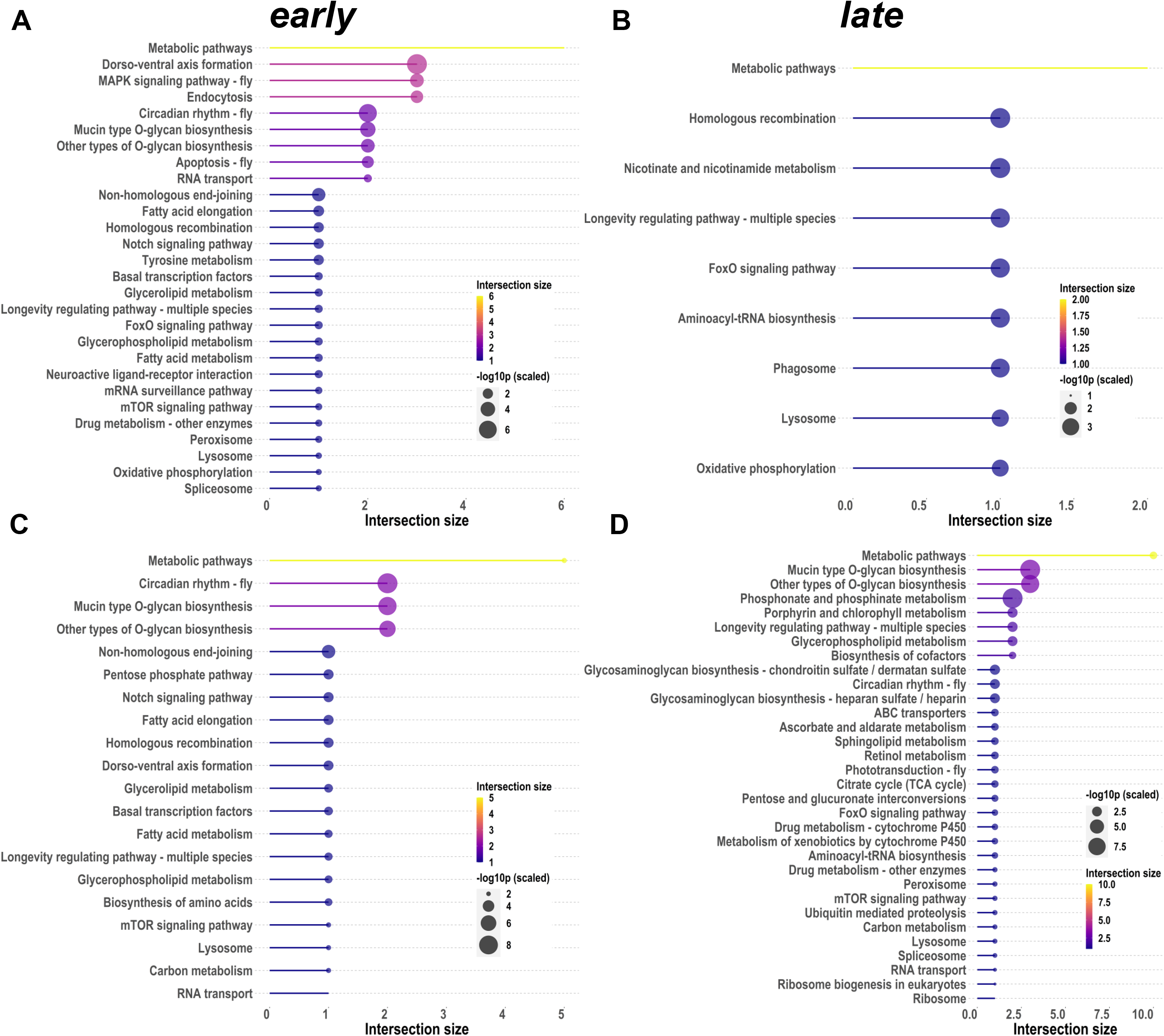
KEGG pathway summaries for genes identified by allele-frequency comparisons. (A, B) Pathways represented in gene sets from control-early (A) and control-late (B) FST comparisons. (C, D) Corresponding pathway summaries for CMH comparisons. Horizontal position and line length indicate intersection size, the number of submitted genes assigned to a pathway; color also encodes intersection size. Bubble size represents -log10(p), scaled for display, from the g:Profiler analysis. The plots are descriptive summaries of the supplied pathway results; display of a pathway does not by itself establish statistical significance after multiple-testing correction.

**Table 3.**
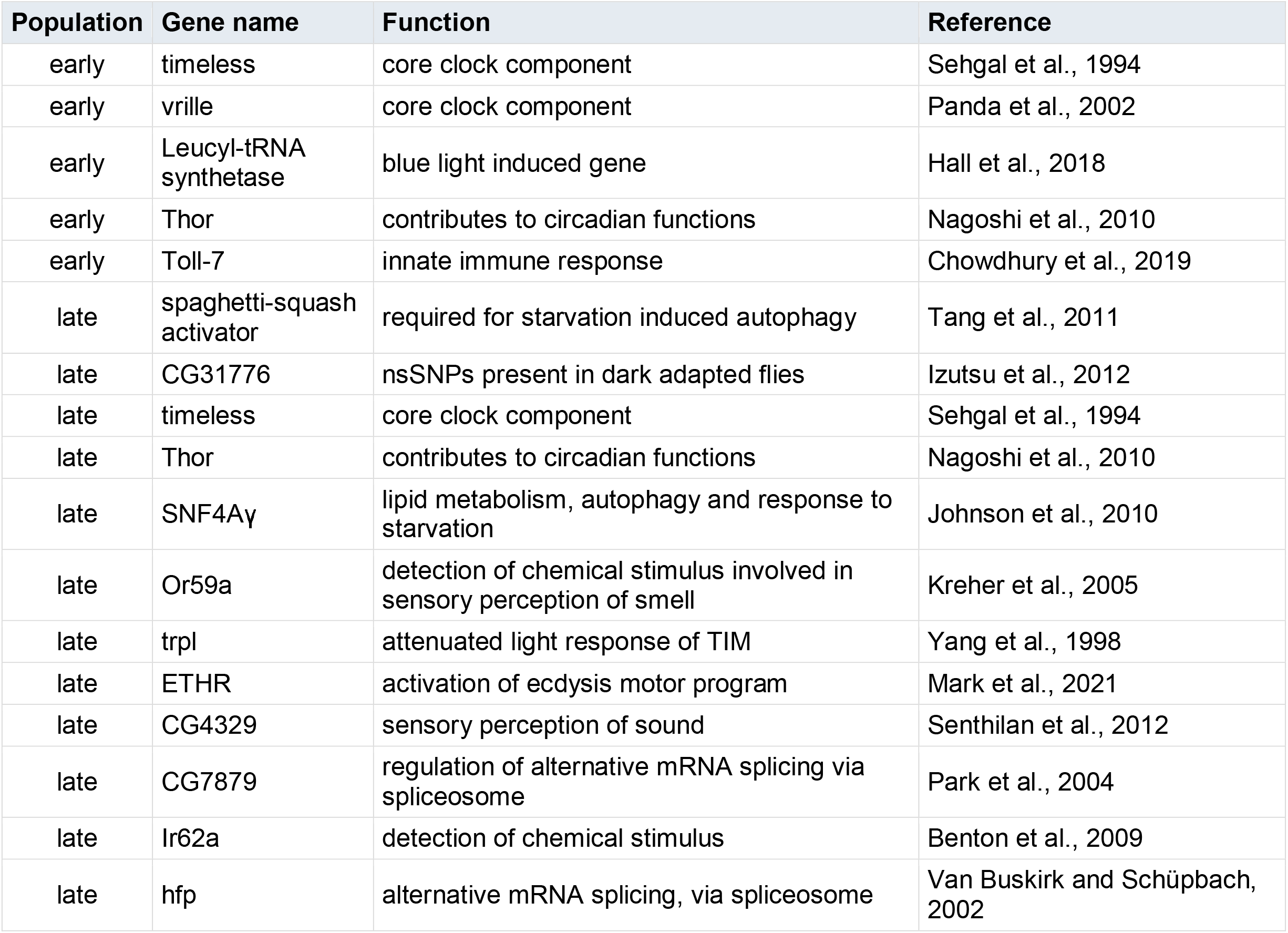
Selected genes identified by CMH comparisons. Selected genes associated with SNPs from the early and late CMH candidate sets. The early threshold was derived by permutation; the late threshold was relaxed to select the top 333 SNPs. Functions and references describe reported biological roles.

### Candidate selective-sweep regions

Pool-hmm detected sweep regions in all three selection regimes (Fig. S7G–I). Regions shared among all four replicate populations spanned approximately 2.42 Mb in *controls*, 1.02 Mb in early populations and 2.90 Mb in *late* populations. After subtracting *control* sweep regions, approximately 0.41 Mb in early populations and 1.34 Mb in *late* populations remained exclusive to the respective selected regime (Fig. S7A–F; Supplementary Tables S7 and S8).

## Discussion

In this study we compared genomes of populations of *Drosophila melanogaster* which evolved significantly divergent chronotypes from their controls using a long-term experimental evolution experiment for about 15 years and ∼260 generations. These populations are unique in terms of their strict maintenance regime, insights provided on the evolution of circadian clock properties with chronotypes, and population level genetically independent replicates. All these characteristics make them ideal candidates for population genomic studies investigating genome-wide changes in evolved divergent chronotypes, in this case, *early* and *late* eclosion rhythm chronotypes. As expected, we observed all populations maintain high level of standing genetic variation (average *D* value > 0; high average/median π value compared to other studies (Yukilevich et al., 2010)), and there were signatures of selection present in the form of selective sweeps (using *Pool-hmm*) in all 12 samples (four replicates each of *early*, *control* and *late* populations) or regions of significantly low *D* value in *early* and *late* populations compared to *control* populations (Fig. 2C,D; Figs. S1 and S7). Though signatures of selection were present in all populations, including *control*, there were large number of loci which were exclusive to *early* and *late* populations compared to *control*. When we investigated allele frequency changes of different SNPs in *early* and *late* populations compared to the *control* population, we found a high number of SNPs where allele frequency had changed (Figs. 2–4). We detected significantly different SNPs through different methods (F_ST_ and CMH) and analyzed each list of SNPs separately as different methodology of detection of allele frequency changes have their own merits and demerits (Figs. 3 and 4) (Kofler, Pandey, et al., 2011; Vlachos et al., 2019). Using these different methods, we identified different sets of genes harboring significantly differentiated SNPs in *early* and *late* populations, compared to the *control* population (Figs. 2–5). We also identified several enriched pathways plausibly under differential selection in these two different populations.

### Photosensitivity, sleep, and clock related changes in *early* populations

In the *early* populations, genes with significantly differentiated SNPs included light-mediated (induced), photoreceptor regulation, and phototransduction genes like *nord*, *Arrestin 1*, *reduce ocelli*, *ninaD*, *notch*, *Basigin*, and *Leucyl-tRNA synthetase* (summarized in *Tables 1–3*) (Caldwell et al., 2007; Damulewicz et al., 2019; Gu et al., 2004; Hall et al., 2018; Kahali et al., 2009; Munro et al., 2010). Previous studies on these populations have shown *early* populations may have evolved heightened light sensitivity and light-mediated positive masking response, suggesting that there may be photosensitivity differences of the circadian clock between *early* and *late* populations (Ghosh et al., 2021; Kumar et al., 2007; Vaze et al., 2012b). Most of the SNPs present in these genes are in regulatory regions, suggesting that regulatory changes in multiple genes related to light input pathways to the circadian clock can modulate chronotype in *Drosophila melanogaster* (Figs. 3–5). We also identified multiple core circadian clock genes (*timeless*, *vrille, sgg*) and genes affecting circadian functions in *Drosophila* (*nervy*, *Thor*, *spaghetti*, 5-HT1B, and *Fbxl4*) harboring many significantly differentiated SNPs in *early* populations (Fig. 5) (Duvall and Taghert, 2012; Hermann-Luibl et al., 2014; Li et al., 2017; Martinek et al., 2001; Means et al., 2015; Nagoshi et al., 2010; Yuan et al., 2005). These genes can modulate circadian clock function directly or indirectly, and fine tuning of circadian phase may be achieved through variants with predicted molecular consequences annotated by SnpEff (Cingolani et al., 2018). Interestingly, we also find multiple sleep regulating genes having multiple SNPs present in them in *early* populations – *kirre*, *Egfr*, *Tyramine* β *hydroxylase*, and *Nlg4* (Fig. 5) (Crocker and Sehgal, 2008; Dreyer et al., 2019; Foltenyi et al., 2007; Harbison et al., 2017; Li et al., 2013). As sleep and circadian clock are tightly correlated processes, we suggest that along with divergent chronotypes, there would be differences in sleep among the *early* and *late* populations which may be governed by differences in the mentioned genes. Indeed, we find *early* and *late* populations have significantly lower nighttime sleep amount and quality compared to the *control* populations under different light intensities and shorter daylengths (Arijit Ghosh; unpublished data). Directly correlated to the adult eclosion rhythm phenotypes of these populations, we find changes in genes (genes accumulating more SNPs) regulating the ecdysis behavior, like *CCAP-R* and *Eip93F* (Cazzamali et al., 2003; FlyBase). *CCAP-R* is directly involved in the modulation of circadian control of the adult eclosion behavior as shown by abnormal timing of eclosion in populations of flies lacking *CCAP* neurons (Park et al., 2003). *Eip93F* has a tightly regulated temporal expression pattern induced by the ecdysone hormone – only during early stages of *Drosophila* metamorphosis, and *Eip93F* mutants die during early stages of pupal development (Baehrecke and Thummel, 1995; Lee et al., 2000). There were also different genes having significant allele frequency changes related to other biological functions – odorant reception (*Ir64a*), response to auditory stimulus and startle response (*nompC*), synapse organization (*dpr8*), neurotransmission (*pickpocket 29*) and adult locomotory behavior (*bruchpilot*) (Ai et al., 2013; Boyd-Gibbins et al., 2021; Carrillo et al., 2015; Hill et al., 2017; Wagh et al., 2006). In the *early* populations, we see putative biological pathways under selection to be – “circadian rhythm – fly”, “notch signaling pathway”, “spliceosome”, etc., and GO terms represented in the summaries included – “RNA polymerase III type 2 promoter sequence-specific DNA binding”, “transcription factor binding”, “nervous system processes”, “instar larval or pupal development”, “metamorphosis”, etc. Many of these pathways can affect development time, morphogenesis (particularly eclosion, and possibly timing of adult eclosion), expression of circadian genes at specific times directly and indirectly (Fig. 6A,C; Fig. S3A,C). For example, the notch signaling pathway has been implicated to be involved in regulation of sensitivity to sleep loss, which could be one of the pathways by which our *early* population showing less nighttime sleep amount and quality under different light intensities and shorter daylengths (Arijit Ghosh; unpublished data) (Seugnet et al., 2011). All these changes were specific to the *early* populations. When we group genes under selection from different analysis sets, we observe significant enrichment of protein-protein interaction network than expected (*p* value = 0.0291) for genes from all three sets (Tajima’s *D* based comparison, pairwise F_ST_ comparison, and CMH test) with 314 nodes and 347 edges in the network (*Fig. S6A*). When we only used the genes from the allele frequency-based tests (pairwise F_ST_ comparison and CMH test), we do not see significant enrichment of the interaction network (*p* value = 0.272), but we observed 101 nodes and 41 edges in the network (*Figs. S4A and S8A*).

### Clock, mRNA splicing, and ecdysis related changes in *late* populations

In the *late* populations, genes with significantly differentiated SNPs included core circadian clock genes or genes affecting circadian functions – *timeless*, *Thor*, *Blimp-1*, and *spaghetti* (summarized in *Tables 1–3*). Though we also found multiple significantly differentiated SNPs in *timeless* in *early* populations, there were many SNPs in *timeless* exclusive to *late* population which were at very different regions of the *timeless* locus (Summarized in *Supplementary Tables S5 and S6*). There were multiple genes in *late* populations related to brain development (*unplugged*), compound eye development (*Netrin-A*), autophagy regulation (*spaghetti-squash activator* and *SNF4A*γ), etc. (Akin and Zipursky, 2016; Hill et al., 2017; Hirth et al., 2003; Johnson et al., 2010; Lippai et al., 2008; Tang et al., 2011). We also found SNPs in *trpl*, which is responsible for attenuated light response of TIM (Yang et al., 1998). More importantly, we identify several SNPs in genes responsible for regulation of alternative mRNA splicing via spliceosome (CG7879), alternative mRNA splicing, via spliceosome (*hfp*), and identify GO term “mRNA splice site selection” and enriched pathway “spliceosome” – all of which indicate significant changes in the mRNA splicing machinery which has been proposed to regulate period and phases of the circadian clock of *Drosophila melanogaster* directly or indirectly, along with temperature sensitivity of the circadian clock (Fig. 6B,D; Fig. S3B,D) (Martin Anduaga et al., 2019; Majercak et al., 2004; Park et al., 2004; Shakhmantsir et al., 2018; Van Buskirk and Schüpbach, 2002). Indeed, we observed significantly more phase lability and temperature sensitivity, and longer free-running period in the *late* population compared to *control* and *early* populations under temperature cycles in both eclosion and locomotor activity rhythms (Abhilash et al., 2019, 2020; Nikhil, Abhilash, et al., 2016). The *late* population also had SNPs in genes involved in activation of ecdysis motor program (*ETHR*), detection of chemical stimulus involved in sensory perception of smell (*Or59A*), sensory perception of sound and touch (*Rootletin*), determination of adult lifespan (*Sirt4*), along with GO terms represented in the summaries, including “response to stimulus”, “primary metabolic process”, “phagosome”, “lysosome”, “sensory perception of sound”, “sensory perception of mechanical stimulus”, “miRNA loading onto RISC involved in gene silencing by miRNA”, and pathway summaries including “circadian rhythm - fly”, “retinol metabolism”, “phototransduction - fly”, “ubiquitin mediated proteolysis”, “metabolic pathways”, “nicotinate and nicotinamide metabolism”, “longevity regulating pathway – multiple species”, “FoxO signaling pathway” etc. (Fig. 6B,D; Fig. S3B,D) (Chen et al., 2015; Kreher et al., 2005; Mark et al., 2021; Wood et al., 2018). Previously we observed that virgin female flies of the *late* population have lower lifespan compared to the *early* population (Nikhil, Ratna, and Sharma, 2016), the current results lead us to speculate that this may be governed by changes in longevity regulating pathways and determinant of adult lifespan (*Sirt4*). Ecdysis Triggering Hormone Receptor (ETHR) knockdown delays the ecdysis behavior, while overexpression of ETHR can accelerate ecdysis – this suggests that SNPs present within ETHR, may lead to delayed eclosion as observed in the *late* population (Kim et al., 2015). Previous work has shown that FoxO signaling has non cell-autonomous effects on the central circadian clock function, and as *foxo* is expressed majorly in the fat body, it suggests that changes in genes expressed in peripheral tissues can also influence circadian behavior in the *late* population (Zheng et al., 2007). When we group genes under selection from different analysis sets, we observe significant enrichment of protein-protein interaction network than expected (*p* value = 0.00032) for genes from all three sets (Tajima’s *D* based comparison, pairwise F_ST_ comparison, and CMH test) with 214 nodes and 131 edges in the network (*Fig. S6B*). The protein-protein interaction networks were larger in *early* population compared to the *late* population, both in terms of number of nodes and edges – this suggests that in *early* population, the genes that harbor most significantly differentiated SNPs are more intricately connected than in *late* population. When we only used the genes from the allele frequency-based tests (pairwise F_ST_ comparison and CMH test), we again see significant enrichment of the interaction network (*p* value = 0.00376), and we observed 130 nodes and 40 edges in the network (*Figs. S4B and S8B*).

There were also a few genes harboring SNPs different from *control* populations in both *early* and *late* populations. Among these, the most noteworthy is *timeless* which has 14 SNPs common to both *early* and *late* populations, along with SNPs exclusive to both populations. In one study it was shown that increased retention of an alternatively spliced intron in *timeless* leads to delayed accumulation of the TIM protein, which in turn leads to delayed phases of the locomotor activity rhythm under LD12:12 and significantly longer period under DD conditions (Shakhmantsir et al., 2018). Another study identified P-element somatic inhibitor (PSI) to be a regulator of thermosensitive alternative splicing of *timeless*; downregulation of Psi shortens the period of the locomotor activity rhythm and alters the phase under temperature cycles (Foley et al., 2019). Temperature dependent alternative splicing of *timeless* has been postulated to act as a thermometer for the Drosophila circadian clock (Martin Anduaga et al., 2019). Since the first demonstration of a link of alternative splicing of *per* to seasonal adaptation (Majercak et al., 1999), much has been discovered along similar lines – this splicing event can be regulated by photoperiods too along with temperature cycles. Similar studies in other organisms, including plants and fungi have given more confidence on alternative splicing mechanisms affecting overt rhythms controlled by the circadian clock and has been reviewed elsewhere (Bartok et al., 2013). We identify several changes in *timeless* and splicing mechanisms in terms of significant SNPs in our divergent chronotype populations and propose that alternative splicing mechanisms may play a major role in phase and period divergence in divergent chronotypes of *Drosophila melanogaster*. This indicates that *timeless* may have evolved as a major player regulating phase divergence in our populations, along with *vrille* in *early* populations. Apart from these core clock genes, various clock output genes and clock regulatory genes showed significant SNPs exclusively in *early* and *late* populations. Most of the SNPs we detected are in regulatory regions, which correlates well with chronotypes being highly quantitative and polygenic traits. Most of the SNPs we identified may contribute a very small part to the final phase divergence among our populations. As we observed temperature cue dependent phase lability is higher in the *late* populations (Abhilash et al., 2019), we expect this phenomenon to be governed by temperature mediated differential splicing of *timeless* or *period*. This is supported by our finding multiple significant SNPs in *timeless* or different spliceosome complex genes (Martin Anduaga et al., 2019; Majercak et al., 1999, 2004; Shakhmantsir et al., 2018). All the significant SNPs and genes we identified in this study have been selected based on their presence in all four replicate populations, which largely excludes the possibility of them being the result of genetic drift or arising purely because of the selection pressure acting on standing genetic variation. The large number of significant SNPs being present in non-clock genes and the diverse array of non-circadian phenotypes evolving differentially among populations across all four replicates suggest that chronotype divergence in populations does not necessarily stem from changes in clock genes, but can be achieved by multiple small-effect changes in different genes in a population and selection on chronotypes may result in correlated changes in other phenotypes (e.g.: sleep, lifespan, fecundity, non-clock photosensitivity etc.). Our populations were derived from natural populations which we have successfully maintained as large outbreeding populations with large standing genetic variation, hence it is expected that correlation we see among traits or loci will most likely be natural genetic correlation among these traits and loci. Thus, our study addresses a long-standing question in the field.

We identified Toll-7 harboring significantly differentiated SNPs in *early* flies. Toll-7 is characterized in the gene group “Toll-NF-κB Signaling Pathway Core Components” (FBgg0001058) in *Drosophila melanogaster*. The relation between immune system and sleep is ubiquitous, and it has been shown under reduced sleep conditions, the expression of an NF-κB, *Relish* is increased (Williams et al., 2007). We observe significantly lower nighttime sleep in our *early* flies compared to the *control* flies, and this difference in nighttime sleep even increases under different photoperiods. Putting together the decrease in nighttime sleep and changes in immunity related genes, the divergent chronotype flies may show difference in their immune response too, an avenue we are actively pursuing.

Three large-scale GWAS studies conducted by Hu et al., Lane et al., and Jones et al. in 2016 describe SNPs associated with chronotypes in different samples of human populations – Hu et al. on European ancestry individuals from a customer cohort of 23andMe (a personalized genotyping service) where chronotype was only a binary trait, In both Lane et al. and Jones et al, the UK BioBank database was used, which included analyses of more homogenous sample and chronotype outcomes were both binary and continuous traits (Hu et al., 2016; Jones et al., 2016; Lane et al., 2016). All three GWASs identified genes related directly or suspected to influence circadian clock or sleep functions (*PER2*, *RGS16*, *FBXL13*), whereas only one identified gene, *Adenylate kinase 5* (*AK5*) did not have any known link to circadian functions. Genes present in 2 of these GWASs identified *HCRTR2*, *HTR6*, *TNRC6B*, and *APH1A* which were known to either affect sleep/circadian phenotypes or to have a diurnal oscillation themselves; only one gene identified (*ERC2*) by two GWASs did not have any known association with circadian rhythms or sleep (Kalmbach et al., 2017). In contrast, we identified large number of genes with no known association to sleep or circadian phenotypes, harboring multiple low and moderate effect SNPs in divergent chronotype populations, which indicates evolution of a highly intricate and flexible circadian system in *Drosophila melanogaster*.

We did not observe any high effect polymorphism (barring one “stop_gained” variant in the *early*-exclusive CMH list), and majority of the polymorphisms were of “low” and “modifier” effects, and very few of “moderate” effect (*Summarized in Supplementary Tables S1–S4*). This suggests that when selection for chronotype divergence acts on natural, standing genetic variation, multiple small effect variants get differentially selected in different chronotypes, and the abundant high effect genes/polymorphisms observed to be affecting chronotypes in different organisms most probably stem from inbreeding and random fixation of loci in the organisms studied. Overall, we hope this study will serve as an important database of information and enable the identification of putative targets for further work investigating the mechanism by which chronotype divergence may occur in *Drosophila melanogaster*.

## Supplementary figure legends

**Supplementary Figure S1.** Genome-wide nucleotide diversity (π). (A, B) Nucleotide diversity in 50 kb windows advanced in 10 kb steps for control and early (A), and control and late (B), populations. Columns indicate chromosome arms and rows indicate the four replicate populations. Genome-wide median π values were 0.00262 for control, 0.00242 for early, and 0.002641 for late populations. Blue indicates early, green control, and red late populations.

**Supplementary Figure S2.** Genome-wide Watterson’s θW. (A, B) Watterson’s θW in 50 kb windows advanced in 10 kb steps for control and early (A), and control and late (B), populations. Columns indicate chromosome arms and rows indicate the four replicate populations. Genome-wide median θW values were 0.002553 for control, 0.002356 for early, and 0.002578 for late populations. Blue indicates early, green control, and red late populations.

**Supplementary Figure S3.** Gene Ontology summaries and the strict control-late CMH screen. (A, B) Gene Ontology terms represented in genes from control-early (A) and control-late (B) FST comparisons. (C, D) Corresponding summaries for CMH comparisons. Horizontal position and line length show the number of submitted genes assigned to a term (the intersection size). Color represents -log10(p), and bubble size represents the same quantity multiplied by 3.5 for display. Displayed terms include nonsignificant results. (E) Manhattan plot of the control-late CMH comparison supplied for the strict permutation screen, with SNP counts in non-overlapping

0.1 Mb windows beneath the plot and the density scale at right. Horizontal lines and highlighted points are retained from the source plot.

**Supplementary Figure S4.** STRING interaction networks for genes from the FST and CMH comparisons. (A, B) Networks for the control-early (A) and control-late (B) gene sets. Nodes represent protein-coding genes and edges represent STRING interactions; edge color and width reflect the combined interaction score. The complete source networks were reported to contain 101 nodes and 41 edges for early populations and 130 nodes and 40 edges for late populations. Isolated nodes were omitted from these layouts.

**Supplementary Figure S5.** Predicted sequence-consequence composition of exclusive SNP sets. (A, B) Consequences for the 686 early-exclusive (A) and 79 late-exclusive (B) FST SNPs. (C, D) Consequences for the 287 early-exclusive (C) and 287 late-exclusive (D) CMH SNPs after removal of 46 shared SNP positions from the two top-333 sets. Segment labels give the supplied SnpEff consequence category and count. The late CMH set used the relaxed, rank-matched cutoff described in Figure 4.

**Supplementary Figure S6.** STRING interaction networks for genes from the Tajima’s D, FST, and CMH analyses. (A, B) Networks for control-early (A) and control-late (B) gene sets. Nodes represent protein-coding genes and edges represent STRING interactions; edge color and width reflect the combined interaction score. The complete source networks were reported to contain 314 nodes and 347 edges for early populations and 214 nodes and 131 edges for late populations. Isolated nodes were omitted from these layouts.

**Supplementary Figure S7.** Candidate selective-sweep regions identified by Pool-hmm. (A, B, D, E) Regions called in all four early replicates (A, blue), all four control replicates (B and E, green), and all four late replicates (D, orange). (C, F) Early-specific (C) and late-specific (F) regions remaining after subtraction of control regions. The total spans shared across the four replicates within each regime were 1.02 Mb for early, 2.42 Mb for control, and 2.90 Mb for late populations. After control subtraction, the early-specific and late-specific spans were 0.41 Mb and 1.34 Mb, respectively. (G-I) Candidate sweep intervals in each of the four control (G), early (H), and late (I) replicate populations. Chromosome arms and genomic coordinates are shown for each track.

**Supplementary Figure S8.** Alternative orthogonal layouts of the FST-plus-CMH STRING networks. (A, B) The early (A) and late (B) networks shown in Supplementary Figure S4, arranged with orthogonal edges. Nodes represent protein-coding genes and edge color and width reflect the STRING combined interaction score. Isolated nodes were omitted. Complete source-network sizes and definitions are given in Supplementary Figure S4.

Tables 1–3 summarize selected genes identified by three complementary population genomic analyses of early and late Drosophila chronotypes.

## References

Abhilash L and Sharma VK (2016) On the relevance of using laboratory selection to study the adaptive value of circadian clocks. Physiological Entomology. DOI: 10.1111/phen.12158.

Abhilash L, Ghosh A and Sheeba V (2019) Selection for timing of eclosion results in co-evolution of temperature responsiveness in Drosophila melanogaster. Journal of Biological Rhythms 34(6). SAGE Publications Inc.: 1–14. DOI: 10.1177/0748730419877315.

Abhilash L, Kalliyil A and Sheeba V (2020) Responses of activity rhythms to temperature cues evolve in Drosophila populations selected for divergent timing of eclosion. Journal of Experimental Biology 223(11). Company of Biologists Ltd. DOI: 10.1242/jeb.222414.

Aguiar GF, da Silva HP and Marques N (1991) Patterns of daily allocation of sleep periods: a case study in an Amazonian riverine community. Chronobiologia 18(1): 9–19.

Ai M, Blais S, Park JY, et al. (2013) Ionotropic glutamate receptors IR64a and IR8a form a functional odorant receptor complex in Vivo in Drosophila. Journal of Neuroscience 33(26). Society for Neuroscience: 10741–10749. DOI: 10.1523/JNEUROSCI.5419-12.2013.

Akin O and Zipursky SL (2016) Frazzled promotes growth cone attachment at the source of a Netrin gradient in the Drosophila visual system. eLife 5. DOI: 10.7554/eLife.20762.

An H, Zhu Z, Zhou C, et al. (2014) Chronotype and a PERIOD3 variable number tandem repeat polymorphism in Han Chinese pilots. International Journal of Clinical and Experimental Medicine 7(10). E-Century Publishing Corporation: 3770–3776.

Andrews S (2010) FastQC: A quality control tool for high throughput sequence data. Babraham Bioinformatics. Available at: https://www.bioinformatics.babraham.ac.uk/projects/fastqc/.

Archer SN, Robilliard DL, Skene DJ, et al. (2003) A length polymorphism in the circadian clock gene Per3 is linked to delayed sleep phase syndrome and extreme diurnal preference. Sleep 26(4). American Academy of Sleep Medicine: 413–415. DOI: 10.1093/sleep/26.4.413.

Aschoff J and Pohl H (1978) Phase relations between a circadian rhythm and its zeitgeber within the range of entrainment. Die Naturwissenschaften 65(2). Germany: Springer-Verlag: 80–84. DOI: 10.1007/BF00440545.

Aschoff J and Wever R (1962) Über Phasenbeziehungen zwischen biologischer Tagesperiodik und Zeitgeberperiodik. Zeitschrift für Vergleichende Physiologie 46(2). Springer-Verlag: 115–128. DOI: 10.1007/BF00341546.

Baehrecke EH and Thummel CS (1995) The Drosophila E93 Gene from the 93F Early Puff Displays Stage- and Tissue-Specific Regulation by 20-Hydroxyecdysone. Developmental Biology 171(1). Dev Biol: 85–97. DOI: 10.1006/dbio.1995.1262.

Barclay NL, Eley TC, Mill J, et al. (2011) Sleep quality and diurnal preference in a sample of young adults: Associations with 5HTTLPR, PER3, and CLOCK 3111. American Journal of Medical Genetics, Part B: Neuropsychiatric Genetics 156(6): 681–690. DOI: 10.1002/ajmg.b.31210.

Bartok O, Kyriacou CP, Levine J, et al. (2013) Adaptation of molecular circadian clockwork to environmental changes: A role for alternative splicing and miRNAs. Proceedings of the Royal Society B: Biological Sciences. The Royal Society. DOI: 10.1098/rspb.2013.0011.

Biswas S and Akey JM (2006) Genomic insights into positive selection. Trends in Genetics. Elsevier Current Trends. DOI: 10.1016/j.tig.2006.06.005.

Boitard S, Kofler R, Françoise P, et al. (2013) Pool-hmm: A Python program for estimating the allele frequency spectrum and detecting selective sweeps from next generation sequencing of pooled samples. Molecular Ecology Resources 13(2). John Wiley & Sons, Ltd: 337–340. DOI: 10.1111/1755-0998.12063.

Bonduriansky R and Day T (2009) Nongenetic Inheritance and Its Evolutionary Implications. Annual Review of Ecology, Evolution, and Systematics 40(1). Annual Reviews: 103–125. DOI: 10.1146/annurev.ecolsys.39.110707.173441.

Boyd-Gibbins N, Tardieu CH, Blunskyte M, et al. (2021) Turnover and activity-dependent transcriptional control of NompC in the Drosophila ear. iScience 24(5). Elsevier: 102486. DOI: 10.1016/j.isci.2021.102486.

Burke MK, Dunham JP, Shahrestani P, et al. (2010) Genome-wide analysis of a long-term evolution experiment with Drosophila. Nature 467(7315). Nature Publishing Group: 587–590. DOI: 10.1038/nature09352.

Caldwell JC, Fineberg SK and Eberl DF (2007) Reduced ocelli encodes the leucine rich repeat protein pray for elves in Drosophila melanogaster. Fly 1(3). NIH Public Access: 146–152. DOI: 10.4161/fly.4562.

Carpen JD, Archer SN, Skene DJ, et al. (2005) A single-nucleotide polymorphism in the 5′-untranslated region of the hPER2 gene is associated with diurnal preference. Journal of Sleep Research 14(3): 293–297. DOI: 10.1111/j.1365-2869.2005.00471.x.

Carpen JD, Von Schantz M, Smits M, et al. (2006) A silent polymorphism in the PER1 gene associates with extreme diurnal preference in humans. Journal of Human Genetics 51(12): 1122–1125. DOI: 10.1007/s10038-006-0060-y.

Carrillo RA, Özkan E, Menon KP, et al. (2015) Control of Synaptic Connectivity by a Network of Drosophila IgSF Cell Surface Proteins. Cell 163(7). Cell Press: 1770–1782. DOI: 10.1016/j.cell.2015.11.022.

Cassidy JJ, Jha AR, Posadas DM, et al. (2013) MiR-9a minimizes the phenotypic impact of genomic diversity by buffering a transcription factor. Cell. DOI: 10.1016/j.cell.2013.10.057.

Cazzamali G, Hauser F, Kobberup S, et al. (2003) Molecular identification of a Drosophila G protein-coupled receptor specific for crustacean cardioactive peptide. Biochemical and Biophysical Research Communications 303(1). Academic Press: 146–152. DOI: 10.1016/S0006-291X(03)00302-4.

Chen J V., Kao LR, Jana SC, et al. (2015) Rootletin organizes the ciliary rootlet to achieve neuron sensory function in Drosophila. Journal of Cell Biology 211(2). The Rockefeller University Press: 435–453. DOI: 10.1083/jcb.201502032.

Cingolani P, Cunningham F, McLaren W and Wang K (2018) Variant annotations in VCF format. Updated January 2018. Available at: https://pcingola.github.io/SnpEff/adds/VCFannotationformat_v1.0.pdf.

Cingolani P, Platts A, Wang LL, et al. (2012) A program for annotating and predicting the effects of single nucleotide polymorphisms, SnpEff: SNPs in the genome of Drosophila melanogaster strain w1118; iso-2; iso-3. *Fly*. DOI: 10.4161/fly.19695.

Crocker A and Sehgal A (2008) Octopamine regulates sleep in Drosophila through protein kinase A-dependent mechanisms. Journal of Neuroscience 28(38). Society for Neuroscience: 9377–9385. DOI: 10.1523/JNEUROSCI.3072-08a.2008.

Croze M, Wollstein A, Božičević V, et al. (2017) A genome-wide scan for genes under balancing selection in Drosophila melanogaster. BMC Evolutionary Biology 17(1). BioMed Central Ltd.: 1–12. DOI: 10.1186/s12862-016-0857-z.

Damulewicz M, Świątek M, Łoboda A, et al. (2019) Daily regulation of phototransduction, circadian clock, DNA repair, and immune gene expression by heme oxygenase in the retina of drosophila. Genes 10(1). Multidisciplinary Digital Publishing Institute (MDPI). DOI: 10.3390/genes10010006.

Dominoni DM, Helm B, Lehmann M, et al. (2013) Clocks for the city: Circadian differences between forest and city songbirds. Proceedings of the Royal Society B: Biological Sciences 280(1763). Royal Society. DOI: 10.1098/rspb.2013.0593.

Doncheva NT, Morris JH, Gorodkin J, et al. (2019) Cytoscape StringApp: Network Analysis and Visualization of Proteomics Data. Journal of Proteome Research 18(2). J Proteome Res: 623–632. DOI: 10.1021/acs.jproteome.8b00702.

Dowle M and Srinivasan A (2020) data.table: Extension of ‘data.fram’. Available at: https://cran.r-project.org/package=data.table.

Drake CL, Belcher R, Howard R, et al. (2015) Length polymorphism in the Period 3 gene is associated with sleepiness and maladaptive circadian phase in night-shift workers. Journal of Sleep Research 24(3). Blackwell Publishing Ltd: 254–261. DOI: 10.1111/jsr.12264.

Dreyer AP, Martin MM, Fulgham C V., et al. (2019) A circadian output center controlling feeding: Fasting rhythms in Drosophila. PLoS Genetics 15(11). Public Library of Science: e1008478. DOI: 10.1371/journal.pgen.1008478.

Duffy JF and Wright KP (2005) Entrainment of the human circadian system by light. Journal of Biological Rhythms. DOI: 10.1177/0748730405277983.

Duffy JF, Rimmer DW and Czeisler CA (2001) Association of intrinsic circadian period with morningness-eveningness, usual wake time, and circadian phase. Behavioral Neuroscience 115(4): 895–899. DOI: 10.1037/0735-7044.115.4.895.

Dunster GP, de la Iglesia L, Ben-Hamo M, et al. (2018) Sleepmore in Seattle: Later school start times are associated with more sleep and better performance in high school students. Science Advances 4(12). American Association for the Advancement of Science: 6200–6212. DOI: 10.1126/sciadv.aau6200.

Duvall LB and Taghert PH (2012) The circadian neuropeptide pdf signals preferentially through a specific adenylate cyclase isoform ac3 in m pacemakers of drosophila. PLoS Biology 10(6). PLoS Biol. DOI: 10.1371/journal.pbio.1001337.

Ebisawa T, Uchiyama M, Kajimura N, et al. (2001) Association of structural polymorphisms in the human period3 gene with delayed sleep phase syndrome. EMBO Reports 2(4): 342–346. DOI: 10.1093/embo-reports/kve070.

Fabian DK, Kapun M, Nolte V, et al. (2012) Genome-wide patterns of latitudinal differentiation among populations of Drosophila melanogaster from North America. Molecular Ecology 21(19). Mol Ecol: 4748–4769. DOI: 10.1111/j.1365-294X.2012.05731.x.

Foley LE, Ling J, Joshi R, et al. (2019) Drosophila psi controls circadian period and the phase of circadian behavior under temperature cycle via tim splicing. eLife 8. eLife Sciences Publications Ltd. DOI: 10.7554/eLife.50063.

Foltenyi K, Greenspan RJ and Newport JW (2007) Activation of EGFR and ERK by rhomboid signaling regulates the consolidation and maintenance of sleep in Drosophila. Nature Neuroscience 10(9). Nat Neurosci: 1160–1167. DOI: 10.1038/nn1957.

Frías-Lasserre D, S AL and Villagra CA (2019) Differences in larval emergence chronotypes for sympatric Rhagoletis brncici Frías and Rhagoletis conversa (Bréthes) (Diptera, Tephritidae). Revista Brasileira de Entomologia 63(3). Sociedade Brasileira De Entomologia: 195–198. DOI: 10.1016/j.rbe.2019.04.003.

Futschik A and Schlötterer C (2010) The next generation of molecular markers from massively parallel sequencing of pooled DNA samples. Genetics 186(1): 207–218. DOI: 10.1534/genetics.110.114397.

García-Allegue R, Lax P, Madariaga AM, et al. (1999) Locomotor and feeding activity rhythms in a light-entrained diurnal rodent, Octodon degus. American Journal of Physiology - Regulatory Integrative and Comparative Physiology 277(2 46-2). DOI: 10.1152/ajpregu.1999.277.2.r523.

Garland T Jr and Rose MR (eds) (2009) Experimental Evolution: Concepts, Methods, and Applications of Selection Experiments. University of California Press. Available at: https://www.ucpress.edu/books/experimental-evolution/pdf.

Gel B (2020) karyoploteR: Plot customizable linear genomes displaying arbitrary data. Available at: https://github.com/bernatgel/karyoploteR.

Gel B and Serra E (2017) karyoploteR: an R / Bioconductor package to plot customizable genomes displaying arbitrary data. Bioinformatics 33(19): 3088–3090. DOI: 10.1093/bioinformatics/btx346.

Ghosh A, Sharma P, Dansana S, et al. (2021) Evidence for Co-Evolution of Masking With Circadian Phase in Drosophila Melanogaster. Journal of Biological Rhythms 36(3). SAGE PublicationsSage CA: Los Angeles, CA: 254–270. DOI: 10.1177/0748730421997262.

Giannotti F, Cortesi F, Sebastiani T, et al. (2002) Circadian preference, sleep and daytime behaviour in adolescence. Journal of Sleep Research 11(3). J Sleep Res: 191–199. DOI: 10.1046/j.1365-2869.2002.00302.x.

Goldin AP, Sigman M, Braier G, et al. (2020) Interplay of chronotype and school timing predicts school performance. Nature Human Behaviour 4(4). Nature Publishing Group: 387–396. DOI: 10.1038/s41562-020-0820-2.

Gottlieb DJ, O’Connor GT, Wilk JB, et al. (2007) Genome-wide association of sleep and circadian phenotypes. BMC Med Genet. DOI: 10.1186/1471-2350-8-S1-S9.

Graves JLL, Hertweck KLL, Phillips MAA, et al. (2017) Genomics of parallel experimental evolution in drosophila. Molecular Biology and Evolution 34(4). Oxford Academic: 831–842. DOI: 10.1093/molbev/msw282.

Gu G, Yang J, Mitchell KA, et al. (2004) Drosophila NinaB and NinaD Act Outside of Retina to Produce Rhodopsin Chromophore. Journal of Biological Chemistry 279(18). Elsevier: 18608–18613. DOI: 10.1074/jbc.M400323200.

Hall H, Ma J, Shekhar S, et al. (2018) Blue light induces a neuroprotective gene expression program in Drosophila photoreceptors. BMC Neuroscience 19(1). BioMed Central. DOI: 10.1186/s12868-018-0443-y.

Harbison ST, Serrano Negron YL, Hansen NF, et al. (2017) Selection for long and short sleep duration in Drosophila melanogaster reveals the complex genetic network underlying natural variation in sleep. PLoS Genetics 13(12). Public Library of Science: e1007098. DOI: 10.1371/journal.pgen.1007098.

Hartl DL and Clark AG (1989) Principles of Population Genetics. 2nd ed. Sinauer Associates, Sunderland, MA.

Helm B and Visser ME (2010) Heritable circadian period length in a wild bird population. Proceedings of the Royal Society B: Biological Sciences 277(1698). Royal Society: 3335–3342. DOI: 10.1098/rspb.2010.0871.

Hermann-Luibl C, Yoshii T, Senthilan PR, et al. (2014) The ion transport peptide is a new functional clock neuropeptide in the fruit fly drosophila melanogaster. Journal of Neuroscience 34(29). J Neurosci: 9522–9536. DOI: 10.1523/JNEUROSCI.0111-14.2014.

Hill A, Zheng X, Li X, et al. (2017) The Drosophila postsynaptic DEG/ENaC channel ppk29 contributes to excitatory neurotransmission. Journal of Neuroscience 37(12). Society for Neuroscience: 3171–3180. DOI: 10.1523/JNEUROSCI.3850-16.2017.

Hirth F, Kammermeier L, Frei E, et al. (2003) An urbilaterian origin of the tripartite brain: Developmental genetic insights from Drosophila. *Development*. Development. DOI: 10.1242/dev.00438.

Hu Y, Shmygelska A, Tran D, et al. (2016) GWAS of 89,283 individuals identifies genetic variants associated with self-reporting of being a morning person. Nature Communications 7: 10448. DOI: 10.1038/ncomms10448.

Hur YM, Bouchard TJ and Lykken DT (1998) Genetic and environmental influence on morningness-eveningness2. Personality and Individual Differences 25(5). Elsevier BV: 917–925. DOI: 10.1016/s0191-8869(98)00089-0.

Hutter S, Li H, Beisswanger S, et al. (2007) Distinctly different sex ratios in African and European populations of Drosophila melanogaster inferred from chromosomewide single nucleotide polymorphism data. Genetics 177(1). Genetics: 469–480. DOI: 10.1534/genetics.107.074922.

Iwase T, Kajimura N, Uchiyama M, et al. (2002) Mutation screening of the human Clock gene in circadian rhythm sleep disorders. Psychiatry Research 109(2): 121–128. DOI: 10.1016/S0165-1781(02)00006-9.

Johnson EC, Kazgan N, Bretz CA, et al. (2010) Altered metabolism and persistent starvation behaviors caused by reduced AMPK function in Drosophila. PLoS ONE 5(9). Public Library of Science: 1–11. DOI: 10.1371/journal.pone.0012799.

Jones SE, Lane JM, Wood AR, et al. (2019) Genome-wide association analyses of chronotype in 697,828 individuals provides insights into circadian rhythms. Nature Communications 10(1). Nature Publishing Group. DOI: 10.1038/s41467-018-08259-7.

Jones SE, Tyrrell J, Wood AR, et al. (2016) Genome-Wide Association Analyses in 128,266 Individuals Identifies New Morningness and Sleep Duration Loci. PLoS Genetics 12(8). Public Library of Science. DOI: 10.1371/journal.pgen.1006125.

Juda M, Vetter C and Roenneberg T (2013) Chronotype modulates sleep duration, sleep quality, and social jet lag in shift-workers. Journal of Biological Rhythms 28(2). SAGE PublicationsSage CA: Los Angeles, CA: 141–151. DOI: 10.1177/0748730412475042.

Kahali B, Bose A, Karandikar U, et al. (2009) On the mechanism underlying the divergent retinal and bristle defects of M8* (E(spl)D) in drosophila. Genesis 47(7). John Wiley & Sons, Ltd: 456–468. DOI: 10.1002/dvg.20521.

Kalmbach DA, Schneider LD, Cheung J, et al. (2017) Genetic basis of chronotype in humans: Insights from three landmark gwas. *Sleep*. Associated Professional Sleep Societies,LLC. DOI: 10.1093/sleep/zsw048.

Kapun M, Barron MG, Staubach F, et al. (2020) Genomic analysis of european drosophila melanogaster populations reveals longitudinal structure, continent-wide selection, and previously unknown DNA viruses. Molecular Biology and Evolution 37(9). Oxford Academic: 2661–2678. DOI: 10.1093/molbev/msaa120.

Katzenberg D, Young T, Finn L, et al. (1998) A CLOCK polymorphism associated with human diurnal preference. Sleep 21(6). American Sleep Disorders Association: 569–576. DOI: 10.1093/sleep/21.6.569.

Kawecki TJ, Lenski RE, Ebert D, Hollis B, Olivieri I and Whitlock MC (2012) Experimental evolution. Trends in Ecology & Evolution 27(10): 547–560. DOI: 10.1016/j.tree.2012.06.001.

Kerkhof GA and Van Dongen HPA (1996) Morning-type and evening-type individuals differ in the phase position of their endogenous circadian oscillator. Neuroscience Letters 218(3): 153–156. DOI: 10.1016/S0304-3940(96)13140-2.

Kim DH, Han MR, Lee G, et al. (2015) Rescheduling Behavioral Subunits of a Fixed Action Pattern by Genetic Manipulation of Peptidergic Signaling. PLoS Genetics 11(9). Public Library of Science: e1005513. DOI: 10.1371/journal.pgen.1005513.

Klei L, Reitz P, Miller M, et al. (2005) Heritability of morningness-eveningness and self-report sleep measures in a family-based sample of 521 hutterites. Chronobiology International 22(6): 1041–1054. DOI: 10.1080/07420520500397959.

Koboldt DC, Chen K, Wylie T, et al. (2009) VarScan: Variant detection in massively parallel sequencing of individual and pooled samples. Bioinformatics. DOI: 10.1093/bioinformatics/btp373.

Kofler R, Orozco-terWengel P, de Maio N, et al. (2011) Popoolation: A toolbox for population genetic analysis of next generation sequencing data from pooled individuals. PLoS ONE 6(1). Public Library of Science: e15925. DOI: 10.1371/journal.pone.0015925.

Kofler R, Pandey RV and Schlötterer C (2011) PoPoolation2: Identifying differentiation between populations using sequencing of pooled DNA samples (Pool-Seq). Bioinformatics. DOI: 10.1093/bioinformatics/btr589.

Koike N, Yoo SH, Huang HC, et al. (2012) Transcriptional architecture and chromatin landscape of the core circadian clock in mammals. Science 338(6105). Science: 349–354. DOI: 10.1126/science.1226339.

Koskenvuo M, Hublin C, Partinen M, et al. (2007) Heritability of diurnal type: A nationwide study of 8753 adult twin pairs. Journal of Sleep Research 16(2): 156–162. DOI: 10.1111/j.1365-2869.2007.00580.x.

Kreher SA, Kwon JY and Carlson JR (2005) The molecular basis of odor coding in the Drosophila larva. Neuron 46(3). Cell Press: 445–456. DOI: 10.1016/j.neuron.2005.04.007.

Kumar S, Kumar D, Paranjpe D a, et al. (2007) Selection on the timing of adult emergence results in altered circadian clocks in fruit flies Drosophila melanogaster. The Journal of experimental biology 210(Pt 5): 906–18. DOI: 10.1242/jeb.001354.

Kunorozva L, Stephenson KJ, Rae DE, et al. (2012) Chronotype and PERIOD3 variable number tandem repeat polymorphism in individual sports athletes. Chronobiology International 29(8): 1004–1010. DOI: 10.3109/07420528.2012.719966.

Lane JM, Vlasac I, Anderson SG, et al. (2016) Genome-wide association analysis identifies novel loci for chronotype in 100,420 individuals from the UK Biobank. Nature Communications 7: 10889. DOI: 10.1038/ncomms10889.

Langmead B and Salzberg SL (2012) Fast gapped-read alignment with Bowtie 2. Nature Methods 9(4): 357–359. DOI: 10.1038/nmeth.1923.

Lee CY, Wendel DP, Reid P, et al. (2000) E93 directs steroid-triggered programmed cell death in Drosophila. Molecular Cell 6(2). Mol Cell: 433–443. DOI: 10.1016/S1097-2765(00)00042-3.

Lehmann M, Spoelstra K, Visser ME, et al. (2012) Effects of temperature on circadian clock and chronotype: An experimental study on a passerine bird. Chronobiology International 29(8): 1062– 1071. DOI: 10.3109/07420528.2012.707159.

Li H, Handsaker B, Wysoker A, et al. (2009) The Sequence Alignment/Map format and SAMtools. Bioinformatics. DOI: 10.1093/bioinformatics/btp352.

Li Q, Li Y, Wang X, et al. (2017) Fbxl4 Serves as a Clock Output Molecule that Regulates Sleep through Promotion of Rhythmic Degradation of the GABAA Receptor. Current Biology 27(23). Curr Biol: 3616-3625.e5. DOI: 10.1016/j.cub.2017.10.052.

Li Y, Zhou Z, Zhang X, et al. (2013) Drosophila neuroligin 4 regulates sleep through modulating GABA transmission. Journal of Neuroscience 33(39). Society for Neuroscience: 15545–15554. DOI: 10.1523/JNEUROSCI.0819-13.2013.

LiLin-Yin (2020) CMplot: Circle Manhattan Plot. Available at: https://github.com/YinLiLin/CMplot.

Lippai M, Csikós G, Maróy P, et al. (2008) SNF4Aγ, the Drosophila AMPK γ subunit is required for regulation of developmental and stress-induced autophagy. Autophagy 4(4). Autophagy: 476–486. DOI: 10.4161/auto.5719.

Majercak J, Chen W-F and Edery I (2004) Splicing of the period Gene 3’-Terminal Intron Is Regulated by Light, Circadian Clock Factors, and Phospholipase C. Molecular and Cellular Biology 24(8): 3359– 3372. DOI: 10.1128/mcb.24.8.3359-3372.2004.

Majercak J, Sidote D, Hardin PE, et al. (1999) How a circadian clock adapts to seasonal decreases in temperature and day length. Neuron 24(1): 219–230. DOI: 10.1016/S0896-6273(00)80834-X.

Mark B, Bustos-González L, Cascallares G, et al. (2021) The circadian clock gates Drosophila adult emergence by controlling the timecourse of metamorphosis. Proceedings of the National Academy of Sciences of the United States of America 118(27). National Academy of Sciences. DOI: 10.1073/pnas.2023249118.

Martin Anduaga A, Evantal N, Patop IL, Bartok O, Weiss R and Kadener S (2019) Thermosensitive alternative splicing senses and mediates temperature adaptation in Drosophila. eLife 8: e44642. DOI: 10.7554/eLife.44642.

Martin M (2011) Cutadapt removes adapter sequences from high-throughput sequencing reads. EMBnet.journal 17(1). EMBnet Stichting: 10. DOI: 10.14806/ej.17.1.200.

Martinek S, Inonog S, Manoukian AS, et al. (2001) A role for the segment polarity gene shaggy/GSK-3 in the Drosophila circadian clock. Cell 105(6). Cell Press: 769–779. DOI: 10.1016/S0092-8674(01)00383-X.

Maukonen M, Kanerva N, Partonen T, et al. (2016) The associations between chronotype, a healthy diet and obesity. 33(8). Taylor & Francis: 972–981. DOI: 10.1080/07420528.2016.1183022.

Means JC, Venkatesan A, Gerdes B, et al. (2015) Drosophila Spaghetti and Doubletime Link the Circadian Clock and Light to Caspases, Apoptosis and Tauopathy. PLoS Genetics Taghert PH (ed.) 11(5). Public Library of Science: e1005171. DOI: 10.1371/journal.pgen.1005171.

Mecacci L and Rocchetti G (1998) Morning and evening types: Stress-related personality aspects. Personality and Individual Differences 25(3). Pergamon: 537–542. DOI: 10.1016/s0191-8869(98)00087-7.

Mishima K, Tozawa T, Satoh K, et al. (2005) The 3111T/C polymorphism of hClock is associated with evening preference and delayed sleep timing in a Japanese population sample. American Journal of Medical Genetics - Neuropsychiatric Genetics 133 B(1): 101–104. DOI: 10.1002/ajmg.b.30110.

Munro M, Akkam Y and Curtin KD (2010) Mutational analysis of Drosophila basigin function in the visual system. Gene 449(1–2). Elsevier: 50–58. DOI: 10.1016/j.gene.2009.09.004.

Nagoshi E, Sugino K, Kula E, et al. (2010) Dissecting differential gene expression within the circadian neuronal circuit of Drosophila. Nature Neuroscience 13(1). Howard Hughes Medical Institute: 60–68. DOI: 10.1038/nn.2451.

Nikhil KL and Sharma VK (2017) On the origin and implications of circadian timekeeping: An evolutionary perspective. In: Kumar V (ed) Biological Timekeeping: Clocks, Rhythms and Behaviour. Springer, New Delhi, pp. 81–129. DOI: 10.1007/978-81-322-3688-7_5.

Nikhil KL, Abhilash L, Sharma VK, et al. (2016) Molecular correlates of circadian clocks in fruit fly Drosophila melanogaster populations exhibiting early and late emergence chronotypes. Journal of Biological Rhythms 31(2): 125–141. DOI: 10.1177/0748730415627933.

Nikhil KL, Goirik G, Karatgi R, et al. (2014) Role of temperature in mediating morning and evening emergence chronotypes in fruit flies drosophila melanogaster. Journal of Biological Rhythms 29(6): 427–441. DOI: 10.1177/0748730414553797.

Nikhil KL, Ratna K and Sharma VK (2016) Life-history traits of Drosophila melanogaster populations exhibiting early and late eclosion chronotypes. BMC Evolutionary Biology 16: 46. DOI: 10.1186/s12862-016-0622-3.

Nikhil KL, Vaze KM, Ratna K and Sharma VK (2016) Circadian clock properties of fruit flies Drosophila melanogaster exhibiting early and late emergence chronotypes. Chronobiology International 33(1): 22–38. DOI: 10.3109/07420528.2015.1108981.

Ocampo-Garcés A, Mena W, Hernández F, et al. (2006) Circadian chronotypes among wild-captured west Andean octodontids. Biological Research 39(2): 209–220. DOI: 10.4067/S0716-97602006000200003.

Orozco-Terwengel P, Kapun M, Nolte V, et al. (2012) Adaptation of Drosophila to a novel laboratory environment reveals temporally heterogeneous trajectories of selected alleles. Molecular Ecology 21(20): 4931–4941. DOI: 10.1111/j.1365-294X.2012.05673.x.

Osland TM, Bjorvatn B, Steen VM, et al. (2011) Association study of a variable-number tandem repeat polymorphism in the clock gene period3 and chronotype in norwegian university students. Chronobiology International 28(9): 764–770. DOI: 10.3109/07420528.2011.607375.

Park JH, Schroeder AJ, Helfrich-Förster C, et al. (2003) Targeted ablation of CCAP neuropeptide-containing neurons of Drosophila causes specific defects in execution and circadian timing of ecdysis behavior. Development. The Company of Biologists. DOI: 10.1242/dev.00503.

Park JW, Parisky K, Celotto AM, et al. (2004) Identification of alternative splicing regulators by RNA interference in Drosophila. Proceedings of the National Academy of Sciences of the United States of America 101(45). National Academy of Sciences: 15974–15979. DOI: 10.1073/pnas.0407004101.

Parsons MJ, Lester KJ, Barclay NL, et al. (2014) Polymorphisms in the circadian expressed genes PER3 and ARNTL2 are associated with diurnal preference and GNβ3 with sleep measures. Journal of Sleep Research 23(5). Blackwell Publishing Ltd: 595–604. DOI: 10.1111/jsr.12144.

Pedrazzoli M, Louzada FM, Pereira DS, et al. (2007) Clock polymorphisms and circadian rhythms phenotypes in a sample of the Brazilian population. Chronobiology International 24(1): 1–8. DOI: 10.1080/07420520601139789.

Perea CS, Niño CL, López-León S, et al. (2014) Study of a Functional Polymorphism in the PER3 Gene and Diurnal Preference in a Colombian Sample. The Open Neurology Journal 8(1): 7–10. DOI: 10.2174/1874205x01408010007.

Pereira DS, Tufik S, Louzada FM, et al. (2005) Association of the length polymorphism in the human Per3 gene with the delayed sleep-phase syndrome: Does latitude have an influence upon it? Sleep 28(1). Oxford University Press (OUP): 29–32. DOI: 10.1093/sleep/28.1.29.

R Core Team (2020) R: A Language and Environment for Statistical Computing. Vienna, Austria. Available at: https://www.r-project.org/.

Randler C, Faßl C and Kalb N (2017) From Lark to Owl: developmental changes in morningness-eveningness from new-borns to early adulthood. Scientific Reports 7(March). Nature Publishing Group: 45874. DOI: 10.1038/srep45874.

Reed LK, Lee K, Zhang Z, et al. (2014) Systems genomics of metabolic phenotypes in wild-type Drosophila melanogaster. Genetics. DOI: 10.1534/genetics.114.163857.

Refinetti R, Wassmer T, Basu P, et al. (2016) Variability of behavioral chronotypes of 16 mammalian species under controlled conditions. Physiology and Behavior 161. Elsevier Inc.: 53–59. DOI: 10.1016/j.physbeh.2016.04.019.

Reimand J, Arak T, Adler P, et al. (2016) g:Profiler-a web server for functional interpretation of gene lists (2016 update). Nucleic acids research. DOI: 10.1093/nar/gkw199.

Remolina SC, Chang PL, Leips J, Nuzhdin SV and Hughes KA (2012) Genomic basis of aging and life-history evolution in Drosophila melanogaster. Evolution 66(11): 3390–3403. DOI: 10.1111/j.1558-5646.2012.01710.x.

Robilliard DL, Archer SN, Arendt J, et al. (2002) The 3111 Clock gene polymorphism is not associated with sleep and circadian rhythmicity in phenotypically characterized human subjects. Journal of Sleep Research 11(4): 305–312. DOI: 10.1046/j.1365-2869.2002.00320.x.

Schwartz MD and Smale L (2005) Individual differences in rhythms of behavioral sleep and its neural substrates in nile grass rats. Journal of Biological Rhythms 20(6): 526–537. DOI: 10.1177/0748730405280924.

Sehgal A, Price JL, Man B and Young MW (1994) Loss of circadian behavioral rhythms and per RNA oscillations in the Drosophila mutant timeless. Science 263(5153): 1603–1606. DOI: 10.1126/science.8128246.

Seugnet L, Suzuki Y, Merlin G, et al. (2011) Notch signaling modulates sleep homeostasis and learning after sleep deprivation in drosophila. Current Biology 21(10). NIH Public Access: 835–840. DOI: 10.1016/j.cub.2011.04.001.

Shakhmantsir I, Nayak S, Grant GR, et al. (2018) Spliceosome factors target timeless (tim) mRNA to control clock protein accumulation and circadian behavior in Drosophila. eLife 7. eLife Sciences Publications Ltd: 1–27. DOI: 10.7554/elife.39821.

Shannon P, Markiel A, Ozier O, et al. (2003) Cytoscape: A software Environment for integrated models of biomolecular interaction networks. Genome Research 13(11). Cold Spring Harbor Laboratory Press: 2498–2504. DOI: 10.1101/gr.1239303.

Sheeba V, Nihal M, Mathew SJ, et al. (2001) Does the difference in the timing of eclosion of the fruit fly Drosophila melanogaster reflect differences in the circadian organization? Chronobiology International 18(4): 601–612.

Stuber EF, Dingemanse NJ, Kempenaers B, et al. (2015) Sources of intraspecific variation in sleep behaviour of wild great tits. Animal Behaviour 106. Academic Press: 201–221. DOI: 10.1016/j.anbehav.2015.05.025.

Szklarczyk D, Gable AL, Nastou KC, et al. (2021) The STRING database in 2021: Customizable protein-protein networks, and functional characterization of user-uploaded gene/measurement sets. Nucleic Acids Research 49(D1). Nucleic Acids Res: D605–D612. DOI: 10.1093/nar/gkaa1074.

Taillard J, Philip P, Coste O, et al. (2003) The circadian and homeostatic modulation of sleep pressure during wakefulness differs between morning and evening chronotypes. Journal of Sleep Research 12(4). J Sleep Res: 275–282. DOI: 10.1046/j.0962-1105.2003.00369.x.

Tajima F (1989) Statistical method for testing the neutral mutation hypothesis by DNA polymorphism. Genetics 123(3). Oxford Academic: 585–595. DOI: 10.1093/genetics/123.3.585.

Tang HW, Wang YB, Wang SL, et al. (2011) Atg1-mediated myosin II activation regulates autophagosome formation during starvation-induced autophagy. EMBO Journal 30(4). John Wiley & Sons, Ltd: 636–651. DOI: 10.1038/emboj.2010.338.

Taylor BJ, Bowman MA, Brindle A, et al. (2020) Evening chronotype, alcohol use disorder severity, and emotion regulation in college students. 37(12). Taylor & Francis: 1725–1735. DOI: 10.1080/07420528.2020.1800028.

Tobler R, Franssen SU, Kofler R, et al. (2014) Massive habitat-specific genomic response in D. melanogaster populations during experimental evolution in hot and cold environments. Molecular Biology and Evolution 31(2). Oxford Academic: 364–375. DOI: 10.1093/molbev/mst205.

Turner TL and Miller PM (2012) Investigating natural variation in drosophila courtship song by the evolve and resequence approach. Genetics. DOI: 10.1534/genetics.112.139337.

Turner TL, Stewart AD, Fields AT, et al. (2011) Population-based resequencing of experimentally evolved populations reveals the genetic basis of body size variation in Drosophila melanogaster. PLoS Genetics. DOI: 10.1371/journal.pgen.1001336.

Van Buskirk C and Schüpbach T (2002) half pint regulates alternative splice site selection in Drosophila. Developmental Cell 2(3). Cell Press: 343–353. DOI: 10.1016/S1534-5807(02)00128-4.

Vaze KM, Kannan NN, Abhilash L and Sharma VK (2012a) Chronotype differences in Drosophila are enhanced by semi-natural conditions. Naturwissenschaften 99(11): 967–971. DOI: 10.1007/s00114-012-0978-1.

Vaze KM, Nikhil KL, Abhilash L and Sharma VK (2012b) Early- and late-emerging Drosophila melanogaster fruit flies differ in their sensitivity to light during morning and evening. Chronobiology International 29(6): 674–682. DOI: 10.3109/07420528.2012.680557.

Viola AU, Archer SN, James LMM, et al. (2007) PER3 Polymorphism Predicts Sleep Structure and Waking Performance. Current Biology 17(7): 613–618. DOI: 10.1016/j.cub.2007.01.073.

Vivanco P, Rol MA and Madrid JA (2010) Pacemaker phase control versus masking by light: Setting the circadian chronotype in dual octodon degus. Chronobiology International 27(7): 1365–1379. DOI: 10.3109/07420528.2010.502984.

Vivanco P, Rol MÁ and Madrid JA (2009) Two steady-entrainment phases and graded masking effects by light generate different circadian chronotypes in Octodon degus. Chronobiology International 26(2): 219–241. DOI: 10.1080/07420520902768203.

Vlachos C, Burny C, Pelizzola M, et al. (2019) Benchmarking software tools for detecting and quantifying selection in evolve and resequencing studies. Genome Biology 20(1). BioMed Central: 1–11. DOI: 10.1186/s13059-019-1770-8.

Volk S, Dyroff J, Georgi K, et al. (1994) Subjective sleepiness and physiological sleep tendency in healthy young morning and evening subjects. Journal of Sleep Research 3(3). J Sleep Res: 138–143. DOI: 10.1111/j.1365-2869.1994.tb00120.x.

Von Schantz M, Taporoski TP, Horimoto ARVR, et al. (2015) Distribution and heritability of diurnal preference (chronotype) in a rural Brazilian family-based cohort, the Baependi study. Scientific Reports 5. Nature Publishing Group. DOI: 10.1038/srep09214.

Wagh DA, Rasse TM, Asan E, et al. (2006) Bruchpilot, a protein with homology to ELKS/CAST, is required for structural integrity and function of synaptic active zones in Drosophila. Neuron 49(6). Cell Press: 833–844. DOI: 10.1016/j.neuron.2006.02.008.

Wang M, Zhao Y and Zhang B (2022) SuperExactTest: Exact Test and Visualization of Multi-Set Intersections. Available at: https://github.com/mw201608/SuperExactTest/.

Wiberg RAW, Gaggiotti OE, Morrissey MB, et al. (2017) Identifying consistent allele frequency differences in studies of stratified populations. Methods in Ecology and Evolution 8(12): 1899–1909. DOI: 10.1111/2041-210X.12810.

Wickham H (2016) Ggplot2: Elegant Graphics for Data Analysis. Springer-Verlag New York. Available at: https://ggplot2.tidyverse.org.

Wickham H, Chang W, Henry L, et al. (2020) ggplot2: Create Elegant Data Visualisations Using the Grammar of Graphics. Available at: https://cran.r-project.org/package=ggplot2.

Williams JA, Sathyanarayanan S, Hendricks JC, et al. (2007) Interaction between sleep and the immune response in Drosophila: A role for the NFκB relish. Sleep 30(4). Oxford Academic: 389–400. DOI: 10.1093/sleep/30.4.389.

Wittmann M, Dinich J, Merrow M, et al. (2006) Social jetlag: Misalignment of biological and social time. In: Chronobiology International, 2006, pp. 497–509. Taylor & Francis. DOI: 10.1080/07420520500545979.

Wood JG, Schwer B, Wickremesinghe PC, et al. (2018) Sirt4 is a mitochondrial regulator of metabolism and lifespan in Drosophila melanogaster. Proceedings of the National Academy of Sciences of the United States of America 115(7). National Academy of Sciences: 1564–1569. DOI: 10.1073/pnas.1720673115.

Yang Z, Emerson M, Su HS, et al. (1998) Response of the timeless protein to light correlates with behavioral entrainment and suggests a nonvisual pathway for circadian photoreception. Neuron 21(1). Cell Press: 215–223. DOI: 10.1016/S0896-6273(00)80528-0.

Yuan Q, Lin F, Zheng X, et al. (2005) Serotonin modulates circadian entrainment in Drosophila. Neuron 47(1). Elsevier: 115–127. DOI: 10.1016/j.neuron.2005.05.027.

Yukilevich R, Turner TL, Aoki F, et al. (2010) Patterns and processes of genome-wide divergence between North American and African Drosophila melanogaster. Genetics 186(1). Oxford University Press: 219–239. DOI: 10.1534/genetics.110.117366.

Zhang Y and Emery P (2013) GW182 Controls Drosophila Circadian Behavior and PDF-Receptor Signaling. Neuron 78(1). Neuron: 152–165. DOI: 10.1016/j.neuron.2013.01.035.

Zheng X, Yang Z, Yue Z, et al. (2007) FOXO and insulin signaling regulate sensitivity of the circadian clock to oxidative stress. Proceedings of the National Academy of Sciences of the United States of America 104(40). National Academy of Sciences: 15899–15904. DOI: 10.1073/pnas.0701599104.

Zhou D, Udpa N, Gersten M, et al. (2011) Experimental selection of hypoxia-tolerant Drosophila melanogaster. Proceedings of the National Academy of Sciences. DOI: 10.1073/pnas.1010643108.

