## Supplementary Figure for "Pooled sequencing reveals patterns of underlying genomic divergence in *Drosophila* chronotypes"

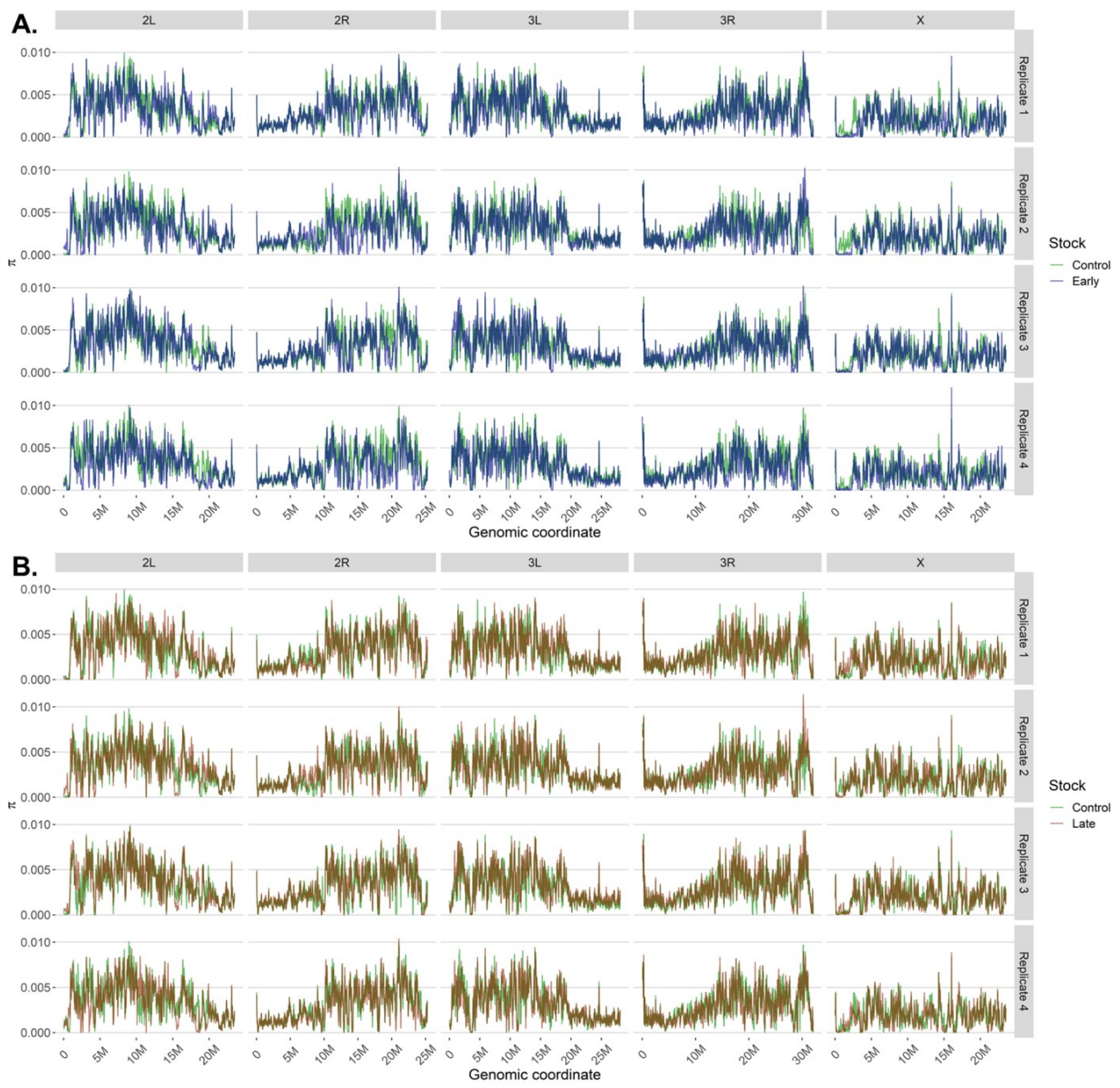


**Supplementary Figure S1.** Genome-wide nucleotide diversity (π). (A, B) Nucleotide diversity in 50 kb windows advanced in 10 kb steps for control and early (A), and control and late (B), populations. Columns indicate chromosome arms and rows indicate the four replicate populations. Genome-wide median π values were 0.00262 for control, 0.00242 for early, and 0.002641 for late populations. Blue indicates early, green control, and red late populations.

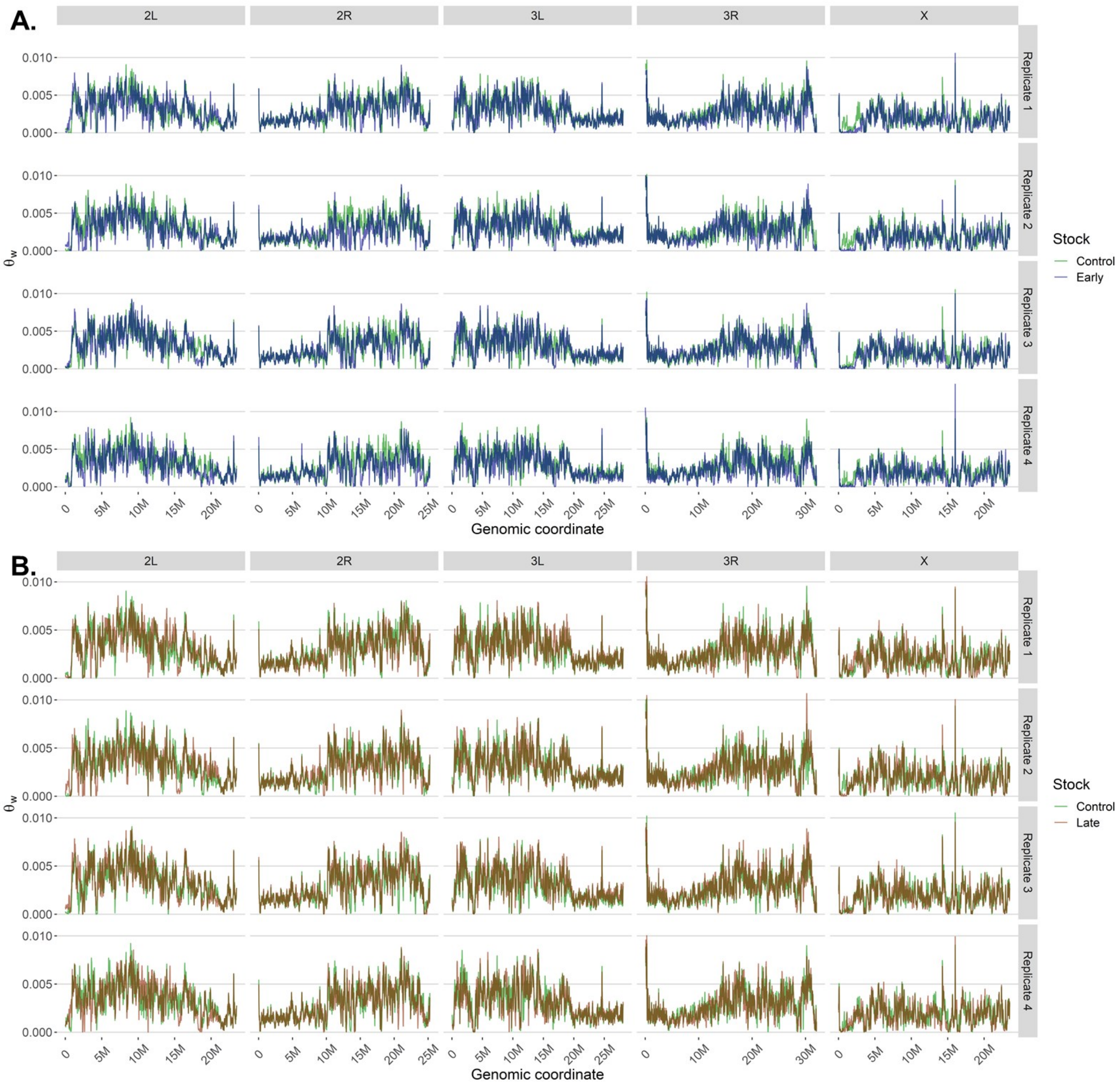


**Supplementary Figure S2.** Genome-wide Watterson's θW. (A, B) Watterson's θW in 50 kb windows advanced in 10 kb steps for control and early (A), and control and late (B), populations. Columns indicate chromosome arms and rows indicate the four replicate populations. Genome-wide median θW values were 0.002553 for control, 0.002356 for early, and 0.002578 for late populations. Blue indicates early, green control, and red late populations.

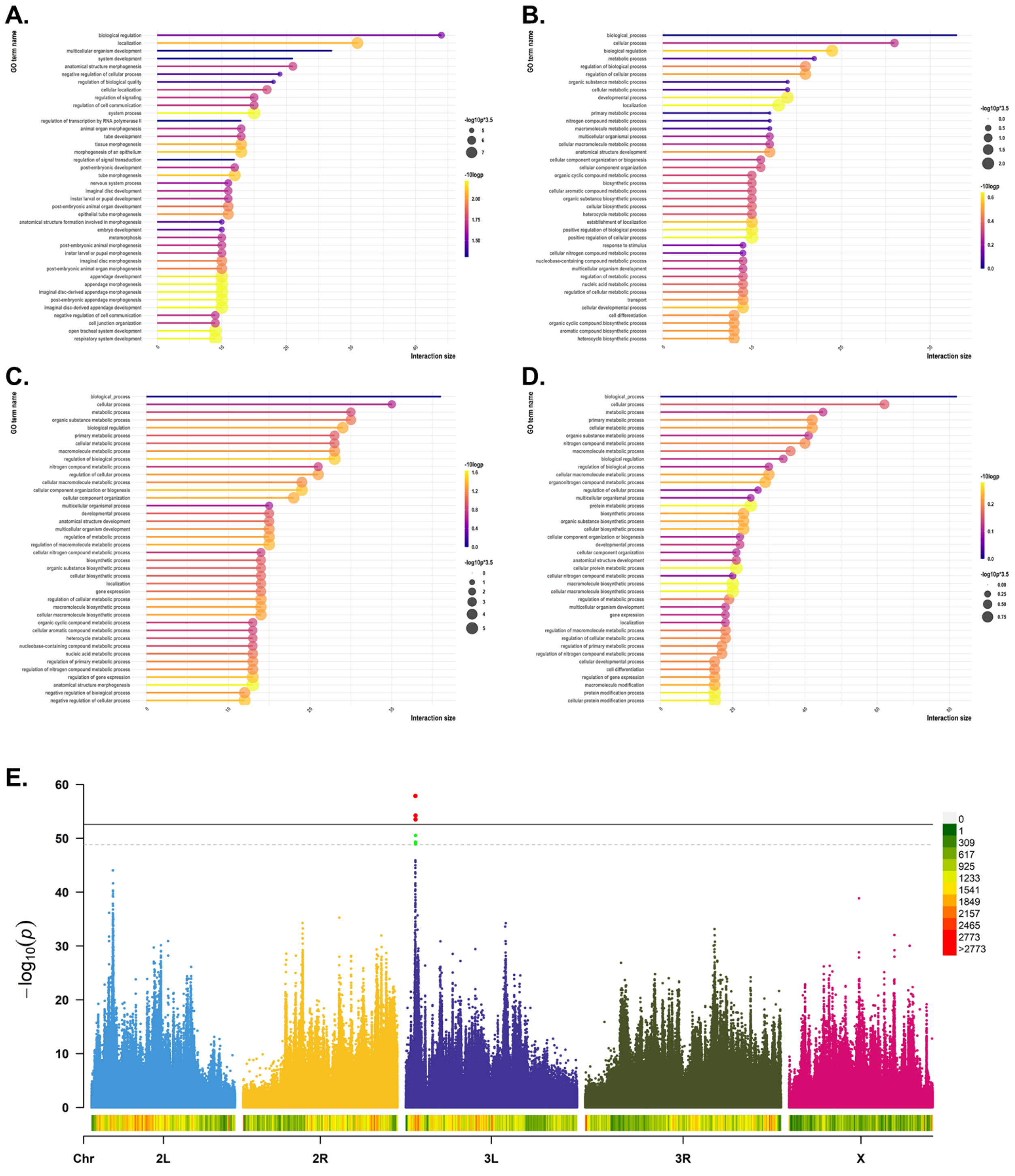
A.
B.
C.
D.
GO term name
Interaction size
-10logp
-log10p*3.5
E.
60
50
40
30
20
10
0
$-\log_{10}(p)$
0
1
309
617
925
1233
1541
1849
2157
2465
2773
>2773
Chr
2L
2R
3L
3R
X

**Supplementary Figure S3.** Gene Ontology summaries and the strict control-late CMH screen. (A, B) Gene Ontology terms represented in genes from control-early (A) and control-late (B) FST comparisons. (C, D) Corresponding summaries for CMH comparisons. Horizontal position and line length show the number of submitted genes assigned to a term (the intersection size). Color represents -log10(p), and bubble size represents the same quantity multiplied by 3.5 for display. Displayed terms include nonsignificant results. (E) Manhattan plot of the control-late CMH comparison supplied for the strict permutation screen, with SNP counts in non-overlapping 0.1 Mb windows beneath the plot and the density scale at right. Horizontal lines and highlighted points are retained from the source plot.

**A.**

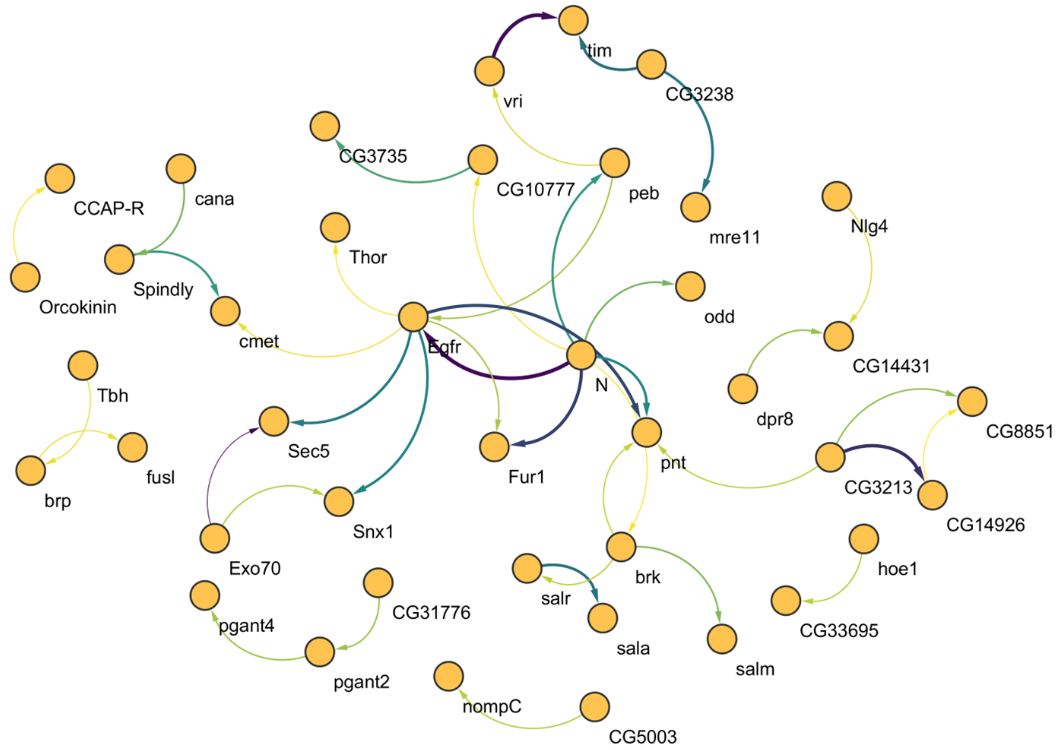
tim
vri
CG3238
CG3735
CG10777
peb
CCAP-R
cana
mre11
Nlg4
Thor
Orcokinin
Spindly
cmet
odd
Egfr
CG14431
Tbh
N
dpr8
CG8851
Sec5
Fur1
pnt
fusl
CG3213
brp
CG14926
Snx1
Exo70
hoe1
brk
salr
CG33695
CG31776
pgant4
sala
salm
pgant2
nompC
CG5003

**B.**

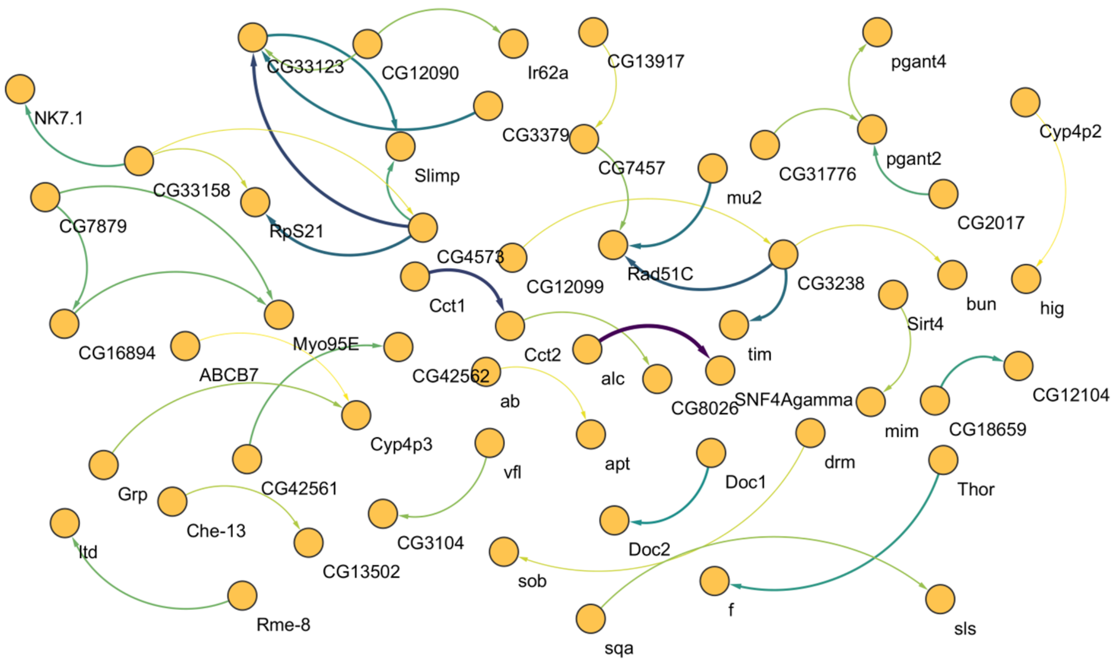
CG33123
CG12090
Ir62a
CG13917
pgant4
NK7.1
CG3379
Cyp4p2
Slimp
CG7457
pgant2
CG33158
mu2
CG31776
CG7879
RpS21
CG2017
CG4573
Rad51C
CG12099
CG3238
bun
hig
Cct1
Sirt4
CG16894
Myo95E
tim
ABCB7
CG42562
Cct2
alc
ab
CG8026
SNF4Agamma
CG12104
mim
CG18659
Cyp4p3
apt
vfl
Grp
CG42561
Doc1
drm
Thor
Che-13
CG3104
Doc2
ltd
CG13502
sob
f
sls
Rme-8
sqa

**Supplementary Figure S4.** STRING interaction networks for genes from the FST and CMH comparisons. (A, B) Networks for the control-early (A) and control-late (B) gene sets. Nodes represent protein-coding genes and edges represent STRING interactions; edge color and width reflect the combined interaction score. The complete source networks were reported to contain 101 nodes and 41 edges for early populations and 130 nodes and 40 edges for late populations. Isolated nodes were omitted from these layouts.

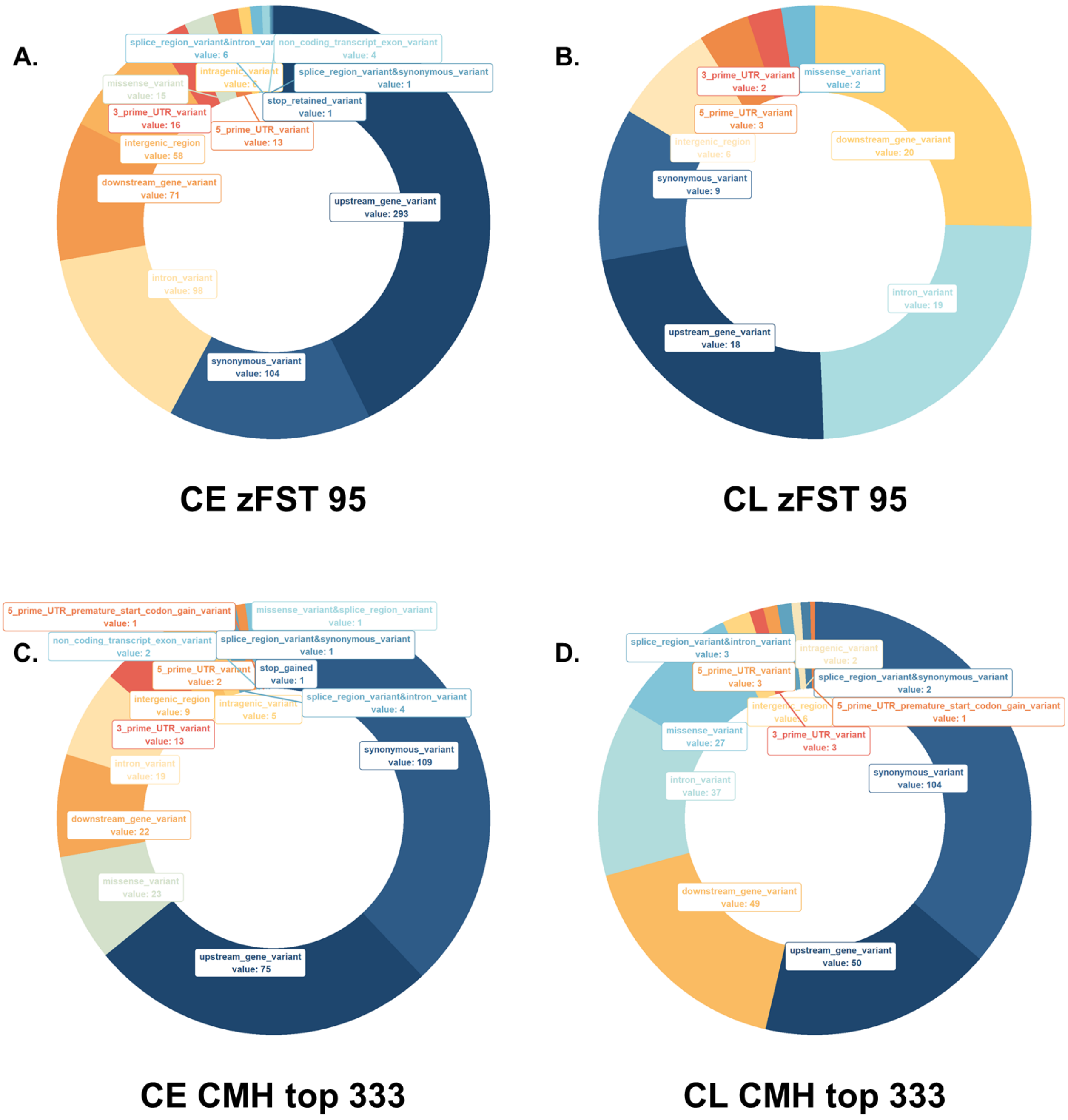


**Supplementary Figure S5.** Predicted sequence-consequence composition of exclusive SNP sets. (A, B) Consequences for the 686 early-exclusive (A) and 79 late-exclusive (B) FST SNPs. (C, D) Consequences for the 287 early-exclusive (C) and 287 late-exclusive (D) CMH SNPs after removal of 46 shared SNP positions from the two top-333 sets. Segment labels give the supplied SnpEff consequence category and count. The late CMH set used the relaxed, rank-matched cutoff described in Figure 4.

**A.**

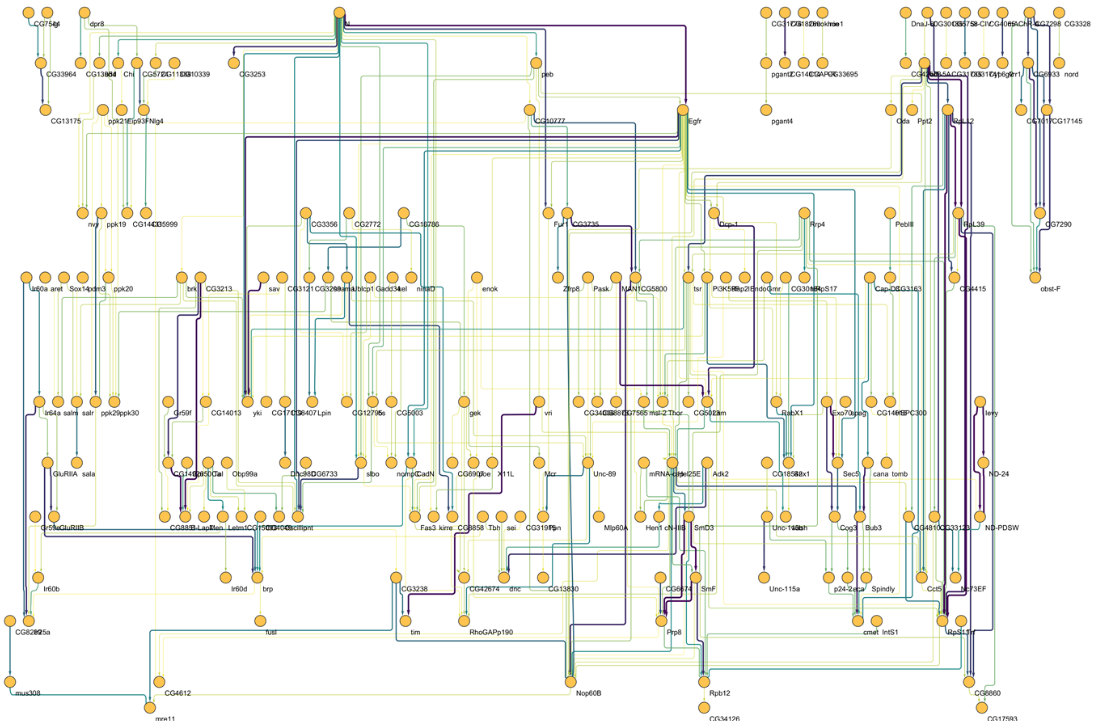

**B.**

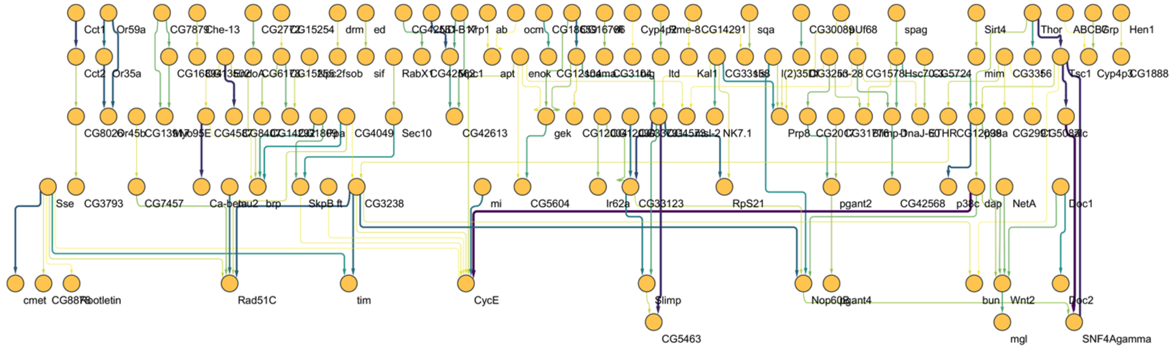

**Supplementary Figure S6.** STRING interaction networks for genes from the Tajima's D, FST, and CMH analyses. (A, B) Networks for control-early (A) and control-late (B) gene sets. Nodes represent protein-coding genes and edges represent STRING interactions; edge color and width reflect the combined interaction score. The complete source networks were reported to contain 314 nodes and 347 edges for early populations and 214 nodes and 131 edges for late populations. Isolated nodes were omitted from these layouts.

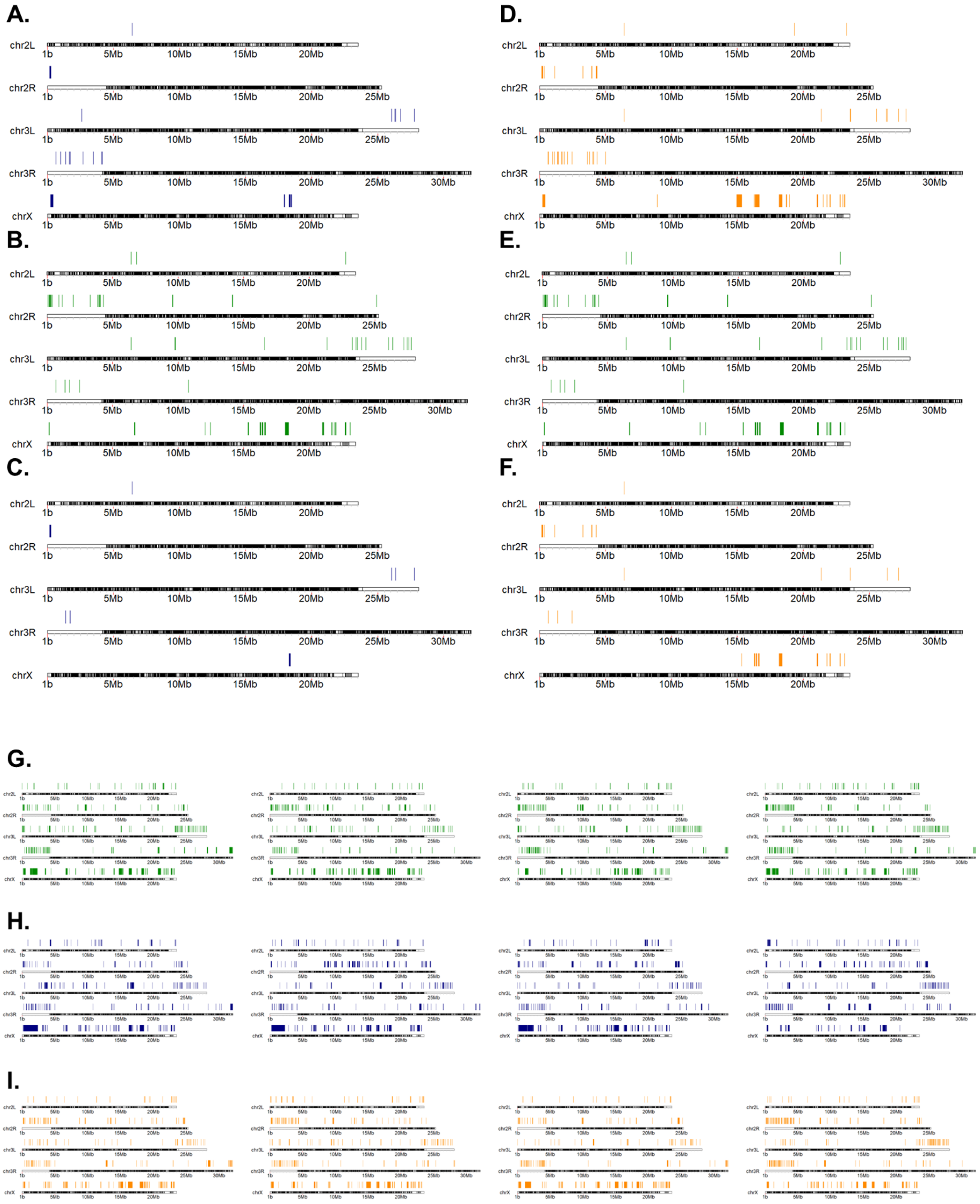
A.
B.
C.
D.
E.
F.
G.
H.
I.
chr2L
chr2R
chr3L
chr3R
chrX
1b
5Mb
10Mb
15Mb
20Mb
25Mb
30Mb

**Supplementary Figure S7.** Candidate selective-sweep regions identified by Pool-hmm. (A, B, D, E) Regions called in all four early replicates (A, blue), all four control replicates (B and E, green), and all four late replicates (D, orange). (C, F) Early-specific (C) and late-specific (F) regions remaining after subtraction of control regions. The total spans shared across the four replicates within each regime were 1.02 Mb for early, 2.42 Mb for control, and 2.90 Mb for late populations. After control subtraction, the early-specific and late-specific spans were 0.41 Mb and 1.34 Mb, respectively. (G-I) Candidate sweep intervals in each of the four control (G), early (H), and late (I) replicate populations. Chromosome arms and genomic coordinates are shown for each track.

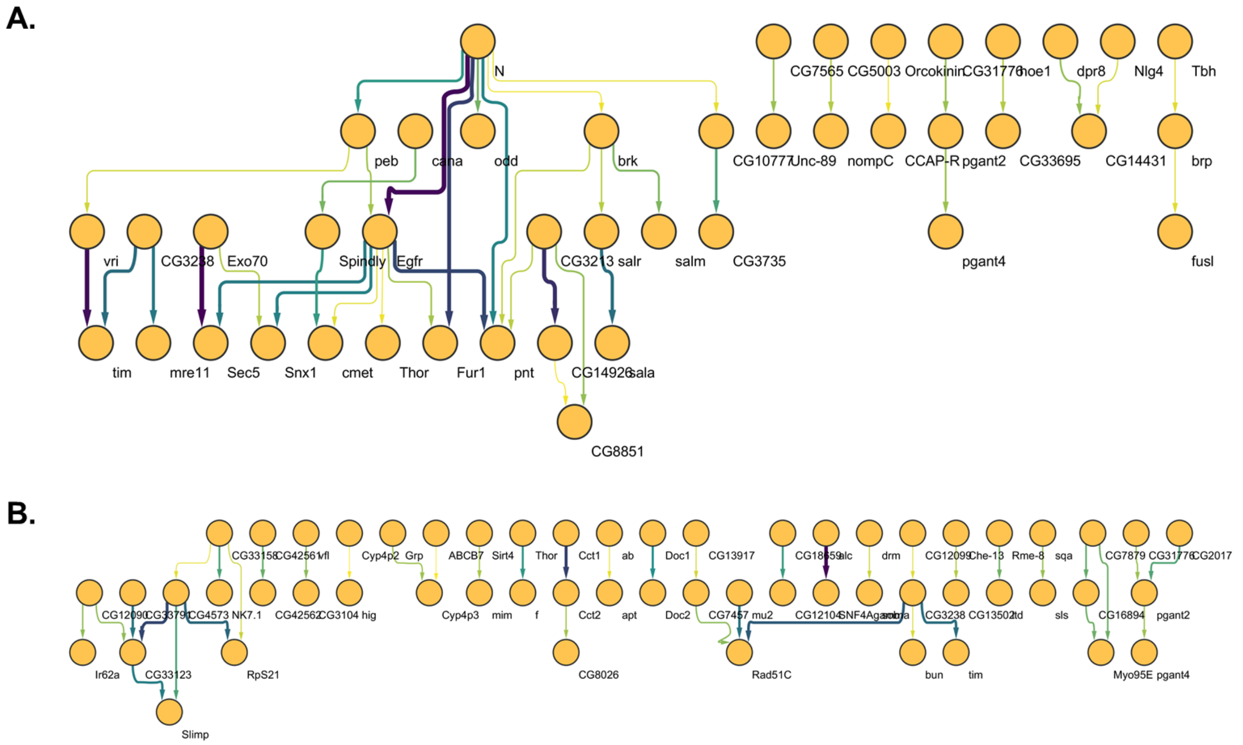


**Supplementary Figure S8.** Alternative orthogonal layouts of the FST-plus-CMH STRING networks. (A, B) The early (A) and late (B) networks shown in Supplementary Figure S4, arranged with orthogonal edges. Nodes represent protein-coding genes and edge color and width reflect the STRING combined interaction score. Isolated nodes were omitted. Complete source-network sizes and definitions are given in Supplementary Figure S4.
