## Supplementary Table for "Pooled sequencing reveals patterns of underlying genomic divergence in *Drosophila* chronotypes"

**Supplementary tables**

Supplementary Tables S1–S9 report variant consequences, clock gene variants, selective sweep intervals, and SNPs shared between early and late CMH candidate sets. SNP, single nucleotide polymorphism; CMH, Cochran–Mantel–Haenszel; UTR, untranslated region.

**Supplementary Table S1. Predicted consequences of early-exclusive FST variants**

Consequences for the 686 early-exclusive SNPs passing the 95th-percentile zFST threshold in all four matched replicate comparisons. Each SNP is represented by its first SnpEff ANN entry. Uppercase labels are impact-category subtotals; indented rows list consequence types within each category.

| **Variant effect or impact category** | **Number of SNPs** |
| --- | --- |
| **LOW** | **112** |
| splice_region_variant&intron_variant | 6 |
| splice_region_variant&synonymous_variant | 1 |
| stop_retained_variant | 1 |
| synonymous_variant | 104 |
| **MODERATE** | **15** |
| missense_variant | 15 |
| **MODIFIER** | **559** |
| 3_prime_UTR_variant | 16 |
| 5_prime_UTR_variant | 13 |
| downstream_gene_variant | 71 |
| intergenic_region | 58 |
| intragenic_variant | 6 |
| intron_variant | 98 |
| non_coding_transcript_exon_variant | 4 |
| upstream_gene_variant | 293 |
| **Grand Total** | **686** |

**Supplementary Table S2. Predicted consequences of late-exclusive FST variants**

Consequences for the 79 late-exclusive SNPs passing the 95th-percentile zFST threshold in all four matched replicate comparisons. Each SNP is represented by its first SnpEff ANN entry. Uppercase labels are impact-category subtotals; indented rows list consequence types within each category.

| **Variant effect or impact category** | **Number of SNPs** |
| --- | --- |
| **LOW** | **9** |
| synonymous_variant | 9 |
| **MODERATE** | **2** |
| missense_variant | 2 |
| **MODIFIER** | **68** |
| 3_prime_UTR_variant | 2 |
| 5_prime_UTR_variant | 3 |
| downstream_gene_variant | 20 |
| intergenic_region | 6 |
| intron_variant | 19 |
| upstream_gene_variant | 18 |
| **Grand Total** | **79** |

**Supplementary Table S3. Predicted consequences of early-exclusive CMH variants**

Consequences for the 287 early-exclusive SNPs after removing the 46 SNPs shared between the two CMH top-333 sets. The early threshold was derived by permutation. Each SNP is represented by its first SnpEff ANN entry. Uppercase labels are impact-category subtotals; indented rows list consequence types within each category.

| **Variant effect or impact category** | **Number of SNPs** |
| --- | --- |
| **HIGH** | **1** |
| stop_gained | 1 |
| **LOW** | **115** |
| 5_prime_UTR_premature_start_codon_gain_variant | 1 |
| splice_region_variant&intron_variant | 4 |
| splice_region_variant&synonymous_variant | 1 |
| synonymous_variant | 109 |
| **MODERATE** | **24** |
| missense_variant | 23 |
| missense_variant&splice_region_variant | 1 |
| **MODIFIER** | **147** |
| 3_prime_UTR_variant | 13 |
| 5_prime_UTR_variant | 2 |
| downstream_gene_variant | 22 |
| intergenic_region | 9 |
| intragenic_variant | 5 |
| intron_variant | 19 |
| non_coding_transcript_exon_variant | 2 |
| upstream_gene_variant | 75 |
| **Grand Total** | **287** |

**Supplementary Table S4. Predicted consequences of late-exclusive CMH variants**

Consequences for the 287 late-exclusive SNPs after removing the 46 SNPs shared between the two CMH top-333 sets. The late threshold was relaxed to select the top 333 SNPs. Each SNP is represented by its first SnpEff ANN entry. Uppercase labels are impact-category subtotals; indented rows list consequence types within each category.

| **Variant effect or impact category** | **Number of SNPs** |
| --- | --- |
| **LOW** | **110** |
| 5_prime_UTR_premature_start_codon_gain_variant | 1 |
| splice_region_variant&intron_variant | 3 |
| splice_region_variant&synonymous_variant | 2 |
| synonymous_variant | 104 |
| **MODERATE** | **27** |
| missense_variant | 27 |
| **MODIFIER** | **150** |
| 3_prime_UTR_variant | 3 |
| 5_prime_UTR_variant | 3 |
| downstream_gene_variant | 49 |
| intergenic_region | 6 |
| intragenic_variant | 2 |
| intron_variant | 37 |
| upstream_gene_variant | 50 |
| **Grand Total** | **287** |

**Supplementary Table S5. Early-exclusive CMH variants in timeless and vrille**

Predicted consequences and impact categories for 15 timeless (tim) and 9 vrille (vri) SNPs in the early-exclusive CMH set. Each row represents one SNP; repeated consequence labels correspond to different SNP records.

| **Type of variant** | **Category of variant** | **Gene name** |
| --- | --- | --- |
| 3_prime_UTR_variant | MODIFIER | tim |
| 3_prime_UTR_variant | MODIFIER | tim |
| synonymous_variant | LOW | tim |
| missense_variant&splice_region_variant | MODERATE | tim |
| intron_variant | MODIFIER | tim |
| intron_variant | MODIFIER | tim |
| intron_variant | MODIFIER | tim |
| intron_variant | MODIFIER | tim |
| intron_variant | MODIFIER | tim |
| synonymous_variant | LOW | tim |
| synonymous_variant | LOW | tim |
| synonymous_variant | LOW | tim |
| 5_prime_UTR_variant | MODIFIER | tim |
| upstream_gene_variant | MODIFIER | tim |
| upstream_gene_variant | MODIFIER | tim |
| upstream_gene_variant | MODIFIER | vri |
| upstream_gene_variant | MODIFIER | vri |
| upstream_gene_variant | MODIFIER | vri |
| upstream_gene_variant | MODIFIER | vri |
| upstream_gene_variant | MODIFIER | vri |
| upstream_gene_variant | MODIFIER | vri |
| upstream_gene_variant | MODIFIER | vri |
| upstream_gene_variant | MODIFIER | vri |
| upstream_gene_variant | MODIFIER | vri |

**Supplementary Table S6. Late-exclusive CMH variants in timeless**

Predicted consequences and impact categories for 9 timeless (tim) SNPs in the late-exclusive CMH set. Each row represents one SNP; repeated consequence labels correspond to different SNP records.

| **Type of variant** | **Category of variant** | **Gene name** |
| --- | --- | --- |
| intron_variant | MODIFIER | tim |
| splice_region_variant&synonymous_variant | LOW | tim |
| synonymous_variant | LOW | tim |
| synonymous_variant | LOW | tim |
| intron_variant | MODIFIER | tim |
| synonymous_variant | LOW | tim |
| synonymous_variant | LOW | tim |
| synonymous_variant | LOW | tim |
| synonymous_variant | LOW | tim |

**Supplementary Table S7. Early-exclusive selective sweep regions and average Tajima’s D**

Genomic intervals identified by Pool-hmm as selective sweeps shared among the four early replicates and exclusive of the control sweep set. The 17 intervals are listed with their average Tajima’s D values.

| **Sweep regions** | **Average D Value** |
| --- | --- |
| 2L:6427152-6443453 | 0.075395422 |
| 2R:165886-241481 | -0.778665112 |
| 2R:243490-286145 | -0.616537239 |
| 3L:24218954-24224496 | -0.356936844 |
| 3L:24595121-24601240 | -0.495584467 |
| 3L:26072652-26089293 | -0.80283812 |
| 3L:26387768-26413463 | -0.710592207 |
| 3L:26836519-26841797 | -0.569878012 |
| 3L:27212412-27222493 | -0.536816679 |
| 3L:27237197-27252886 | -0.571513483 |
| 3L:27565708-27571919 | -0.587773465 |
| 3L:27604816-27609696 | -0.664089313 |
| 3L:27796463-27801898 | -0.679601935 |
| 3R:1378942-1389707 | -0.446114859 |
| 3R:1731842-1750040 | -0.698218262 |
| 3R:1985987-1994833 | -0.435463757 |
| X:18288670-18427409 | -0.879016049 |

**Supplementary Table S8. Late-exclusive selective sweep regions and average Tajima’s D**

Genomic intervals identified by Pool-hmm as selective sweeps shared among the four late replicates and exclusive of the control sweep set. The 53 intervals are listed with their average Tajima’s D values.

| **Sweep regions** | **Average D Value** |
| --- | --- |
| 2L:3203669-3209864 | 0.425528905 |
| 2L:6427152-6444980 | 0.200780694 |
| 2L:13449665-13456771 | 0.593837602 |
| 2L:20215222-20218183 | -0.00836032 |
| 2L:20303232-20312906 | -0.126956155 |
| 2R:167066-286594 | -0.721946371 |
| 2R:406655-430990 | -0.667698296 |
| 2R:663098-668522 | -0.29770442 |
| 2R:699235-703517 | -0.286111925 |
| 2R:1161290-1177433 | -0.581824691 |
| 2R:1234382-1245021 | -0.557363408 |
| 2R:2019474-2030305 | -0.461191546 |
| 2R:2110623-2117977 | -0.35310866 |
| 2R:3039829-3050166 | -0.573716924 |
| 2R:3304209-3313133 | -0.365428129 |
| 2R:3881154-3890194 | -0.457176692 |
| 2R:3960674-3999918 | -0.696019362 |
| 2R:4300402-4307442 | -0.250535576 |
| 2R:4310363-4320760 | -0.616147413 |
| 2R:5902477-5909622 | 0.359176514 |
| 3L:6405972-6431765 | 0.424655068 |
| 3L:21365975-21378683 | -0.056056214 |
| 3L:22990230-22996709 | 0.061662088 |
| 3L:23573960-23578445 | -0.389676089 |
| 3L:26382695-26413457 | -0.584293964 |
| 3L:26674894-26682887 | -0.600560999 |
| 3L:26836552-26844539 | -0.411602348 |
| 3L:27212412-27222542 | -0.490158843 |
| 3L:27225036-27252861 | -0.545141347 |
| 3L:27565631-27571919 | -0.608327628 |
| 3L:27814609-27827044 | -0.514129886 |
| 3R:220082-229674 | -0.464080624 |
| 3R:685326-691751 | -0.648144185 |
| 3R:734298-744666 | -0.479300306 |
| 3R:1369553-1375549 | -0.380533151 |
| 3R:1378915-1389707 | -0.430834124 |
| 3R:1729975-1744282 | -0.751668703 |
| 3R:2477862-2488599 | -0.373920543 |
| 3R:4045480-4057938 | -0.428485674 |
| 3R:7731336-7736692 | 0.383778757 |
| X:1353135-1367904 | -0.397123387 |
| X:15321393-15348579 | -1.010400412 |
| X:16270099-16324145 | -0.732422727 |
| X:16340838-16356930 | -0.212647063 |
| X:16417169-16485959 | -1.071838723 |
| X:16596275-16696614 | -0.639245198 |
| X:18182443-18417500 | -0.763990093 |
| X:21039100-21137905 | -0.241711912 |
| X:21806666-21821848 | -0.366456076 |
| X:22022951-22032923 | -0.398916082 |
| X:22060069-22066818 | -0.378886682 |
| X:22757082-22841415 | -0.682099816 |
| X:23120341-23150558 | -0.635775632 |

**Supplementary Table S9. SNPs shared between the early and late CMH top-333 sets**

The two CMH candidate sets share 46 SNPs assigned to 10 genes or loci by their first SnpEff ANN entry. Panels A and B summarize predicted impacts and primary consequences; panel C lists all 46 SNP records. Coordinates and alleles are reproduced from the source SNP calls. The early threshold was derived by permutation; the late threshold was relaxed to select the top 333 SNPs.

**A Predicted impact**

| **Predicted impact** | **Number of SNPs** |
| --- | --- |
| LOW | 31 |
| MODERATE | 4 |
| MODIFIER | 11 |
| Total | 46 |

**B Primary predicted consequence**

| **Primary predicted consequence** | **Number of SNPs** |
| --- | --- |
| upstream_gene_variant | 4 |
| synonymous_variant | 31 |
| intron_variant | 4 |
| downstream_gene_variant | 1 |
| 3_prime_UTR_variant | 2 |
| missense_variant | 4 |
| Total | 46 |

**C Variant details**

| **Chr** | **Position** | **Ref** | **Alt** | **Primary consequence** | **Impact** | **Gene or locus** |
| --- | --- | --- | --- | --- | --- | --- |
| 2L | 3470962 | A | C | synonymous_variant | LOW | CG31776 |
| 2L | 3470966 | A | C | synonymous_variant | LOW | CG31776 |
| 2L | 3474030 | A | C | synonymous_variant | LOW | pgant4 |
| 2L | 3474349 | C | T | synonymous_variant | LOW | pgant4 |
| 2L | 3475176 | C | T | 3_prime_UTR_variant | MODIFIER | pgant4 |
| 2L | 3481242 | T | C | synonymous_variant | LOW | CG15414 |
| 2L | 3481468 | A | G | missense_variant | MODERATE | CG15414 |
| 2L | 3493336 | C | G | upstream_gene_variant | MODIFIER | CG15414 |
| 2L | 3493361 | G | C | upstream_gene_variant | MODIFIER | CG15414 |
| 2L | 3498274 | G | C | intron_variant | MODIFIER | tim |
| 2L | 3498281 | T | C | intron_variant | MODIFIER | tim |
| 2L | 3499609 | C | T | synonymous_variant | LOW | tim |
| 2L | 3499621 | A | G | synonymous_variant | LOW | tim |
| 2L | 3499666 | T | C | synonymous_variant | LOW | tim |
| 2L | 3501851 | A | T | synonymous_variant | LOW | tim |
| 2L | 3502927 | A | G | synonymous_variant | LOW | tim |
| 2L | 3502978 | C | T | synonymous_variant | LOW | tim |
| 2L | 3503064 | C | T | missense_variant | MODERATE | tim |
| 2L | 3503096 | T | G | missense_variant | MODERATE | tim |
| 2L | 3503140 | T | A | synonymous_variant | LOW | tim |
| 2L | 3503317 | G | A | synonymous_variant | LOW | tim |
| 2L | 3503602 | G | T | synonymous_variant | LOW | tim |
| 2L | 3503924 | T | C | missense_variant | MODERATE | tim |
| 2L | 3511937 | G | T | synonymous_variant | LOW | CG33123 |
| 2L | 3511946 | C | A | synonymous_variant | LOW | CG33123 |
| 2L | 3512033 | G | A | synonymous_variant | LOW | CG33123 |
| 2L | 3513375 | G | A | synonymous_variant | LOW | CG33123 |
| 2L | 3513387 | G | A | synonymous_variant | LOW | CG33123 |
| 2L | 3513652 | G | T | synonymous_variant | LOW | CG33123 |
| 2L | 3513694 | C | T | synonymous_variant | LOW | CG33123 |
| 2L | 3513712 | G | A | synonymous_variant | LOW | CG33123 |
| 2L | 3517706 | A | G | synonymous_variant | LOW | CG3213 |
| 2L | 3518387 | G | A | synonymous_variant | LOW | CG3213 |
| 2L | 3518408 | G | A | synonymous_variant | LOW | CG3213 |
| 2L | 3518429 | T | A | synonymous_variant | LOW | CG3213 |
| 2L | 3519038 | T | C | synonymous_variant | LOW | CG3213 |
| 2L | 3519065 | G | A | synonymous_variant | LOW | CG3213 |
| 2L | 3519146 | A | G | synonymous_variant | LOW | CG3213 |
| 2L | 3519167 | A | G | synonymous_variant | LOW | CG3213 |
| 2L | 3519395 | G | T | synonymous_variant | LOW | CG3213 |
| 2L | 3534447 | C | T | upstream_gene_variant | MODIFIER | Spindly |
| 2L | 3547192 | A | G | intron_variant | MODIFIER | drm |
| 2L | 3547653 | A | G | 3_prime_UTR_variant | MODIFIER | drm |
| 2L | 3548553 | A | G | downstream_gene_variant | MODIFIER | drm |
| 2L | 11344235 | C | A | upstream_gene_variant | MODIFIER | CG14926 |
| 2L | 11365437 | A | G | intron_variant | MODIFIER | salr |
